# A sensorimotor-association axis of thalamocortical connection development

**DOI:** 10.1101/2024.06.13.598749

**Authors:** Valerie J. Sydnor, Joëlle Bagautdinova, Bart Larsen, Michael J. Arcaro, Deanna M. Barch, Dani S. Bassett, Aaron F. Alexander-Bloch, Philip A. Cook, Sydney Covitz, Alexandre R. Franco, Raquel E. Gur, Ruben C. Gur, Allyson P. Mackey, Kahini Mehta, Steven L. Meisler, Michael P. Milham, Tyler M. Moore, Eli J. Müller, David R. Roalf, Taylor Salo, Gabriel Schubiner, Jakob Seidlitz, Russell T. Shinohara, James M. Shine, Fang-Cheng Yeh, Matthew Cieslak, Theodore D. Satterthwaite

**Affiliations:** Penn Lifespan Informatics and Neuroimaging Center (PennLINC), Department of Psychiatry, Perelman School of Medicine, University of Pennsylvania, Philadelphia, PA, USA; Department of Psychiatry, Perelman School of Medicine, University of Pennsylvania, Philadelphia, PA, USA; Department of Psychiatry, University of Pittsburgh Medical Center, University of Pittsburgh, Pittsburgh, PA, USA; Department of Pediatrics, Masonic Institute for the Developing Brain, University of Minnesota, Minneapolis, MN, USA; Department of Psychology, School of Arts and Sciences, University of Pennsylvania, Philadelphia, PA, USA; Department of Psychiatry, Washington University School of Medicine in St Louis, St Louis, Missouri, USA; Department of Psychological & Brain Sciences, Washington University in St Louis, St Louis, Missouri, USA; Department of Bioengineering, University of Pennsylvania, Philadelphia, PA, USA; Department of Electrical & Systems Engineering, University of Pennsylvania, Philadelphia, PA, USA; Department of Physics & Astronomy, University of Pennsylvania, Philadelphia, PA, USA; The Santa Fe Institute, Santa Fe, NM, USA; Lifespan Brain Institute (LiBI), Children’s Hospital of Philadelphia and Penn Medicine, Philadelphia, PA, USA; Department of Child and Adolescent Psychiatry and Behavioral Science, The Children’s Hospital of Philadelphia, Philadelphia, PA, USA; Penn Image Computing and Science Lab (PICSL), Department of Radiology, University of Pennsylvania, Philadelphia, PA, USA; Department of Bioengineering, Schools of Engineering and Medicine, Stanford University, Stanford, CA, USA; Center for Biomedical Imaging and Neuromodulation, Nathan Kline Institute for Psychiatric Research, Orangeburg, NY, USA; Strategic Data Initiatives, Child Mind Institute, New York, NY, USA; Department of Psychiatry, NYU Grossman School of Medicine, New York, NY, USA; Neurodevelopment and Psychosis Section, Department of Psychiatry, Perelman School of Medicine, University of Pennsylvania, Philadelphia, PA, USA; Brain Behavior Laboratory, Department of Psychiatry, Perelman School of Medicine, University of Pennsylvania, Philadelphia, PA, USA; Program in Speech and Hearing Bioscience and Technology, Harvard University, Division of Medical Sciences, Cambridge, MA, USA; Center for the Developing Brain, Child Mind Institute, New York, NY, USA; School of Medical Sciences, Faculty of Medicine and Health, The University of Sydney, Sydney, Australia; Institute for Translational Medicine and Therapeutics, University of Pennsylvania, Philadelphia, PA, USA; Penn Statistics in Imaging and Visualization Center, Department of Biostatistics, Epidemiology, & Informatics, Philadelphia, PA, USA; Center for Biomedical Image Computing and Analytics, Department of Radiology, University of Pennsylvania, Philadelphia, PA, USA; Department of Neurological Surgery, University of Pittsburgh, Pittsburgh, PA, USA

## Abstract

Human cortical development follows a sensorimotor-to-association sequence during childhood and adolescence^1–6^. The brain’s capacity to enact this sequence over decades indicates that it relies on intrinsic mechanisms to regulate inter-regional differences in the timing of cortical maturation, yet regulators of human developmental chronology are not well understood. Given evidence from animal models that thalamic axons modulate windows of cortical plasticity^7–12^, here we evaluate the overarching hypothesis that structural connections between the thalamus and cortex help to coordinate cortical maturational heterochronicity during youth. We first introduce, cortically annotate, and anatomically validate a new atlas of human thalamocortical connections using diffusion tractography. By applying this atlas to three independent youth datasets (ages 8-23 years; total *N* = 2,676), we reproducibly demonstrate that thalamocortical connections develop along a maturational gradient that aligns with the cortex’s sensorimotor-association axis. Associative cortical regions with thalamic connections that take longest to mature exhibit protracted expression of neurochemical, structural, and functional markers indicative of higher circuit plasticity as well as heightened environmental sensitivity. This work highlights a central role for the thalamus in the orchestration of hierarchically organized and environmentally sensitive windows of cortical developmental malleability.

## Introduction

Human cortical development is a protracted process that unfolds in a temporally asynchronous manner, with different cortical regions maturing at different rates. During childhood and adolescence, inter-regional variability in maturational timing (i.e., maturational heterochronicity) is organized by the cortex’s sensorimotor-association (S-A) axis^1^. Accumulating evidence suggests that reductions in plasticity gradually progress with age from the sensorimotor to the association pole of this hierarchical cortical axis^1–6,13–15^. The brain’s capacity to enact an S-A developmental sequence over decades indicates that intrinsic timing mechanisms exist that regulate the relative pace of cortical development in each region. Here, we evaluate the hypothesis that the thalamus and its axonal connections with the cortex play a role in coordinating timescales of cortical developmental plasticity in the human brain. To test this hypothesis non-invasively, we introduce a new diffusion MRI atlas of cortically-annotated thalamocortical structural connections. We then apply this atlas to determine whether connections between the thalamus and cortex exhibit a chronological maturational gradient that aligns with the S-A axis of cortical developmental heterochronicity.

The thalamus is a bilateral gray matter structure in the diencephalon that sends (typically reciprocal) axonal projections throughout the human cortical mantle^16–18^. These direct thalamic projections to cortex are established in prenatal^19^ and early postnatal^20^ development, at which time they begin to have a profound effect on the formation and refinement of cortical circuits. Rodent and non-human primate studies have shown that thalamic inputs to cortex partly control cortical region arealization^21,22^, gene expression^22^, laminar architecture^23^, hierarchical areal identity,^20,22^ and the construction of circuit ensembles^20^. Early cortical sculpting by thalamic inputs has primarily been documented in primary cortices, with recent work extending this developmental phenomenon to association cortex in both mice^24^ and non-human primates^25^. As development progresses, functional interactions between the thalamus and cortex continue to be refined^7,26^ such that by adulthood, thalamocortical pathways gate information transfer along sensorimotor-to-associative cortical hierarchies^18,26–30^. The thalamus therefore helps to govern both spatially local cortical circuit maturation and the emergence of hierarchically organized temporal dynamics—making it well positioned to regulate a spatiotemporal maturational program that progresses along the S-A axis.

Axonal projections from the thalamus to cortex may impact not only cortical area properties during development, but also time windows of cortical developmental plasticity. In the murine brain, periods of heightened experience-dependent cortical plasticity co-occur with the assembly^8^, reorganization^9^, and normative strengthening^7,10,12,31^ of thalamocortical axons. Observed relationships between the expression of cortical plasticity and refinements in thalamocortical connectivity may have origins in thalamic modulation of parvalbumin (PV) positive cortical interneurons—inhibitory cells that receive potent thalamic synapses in development^11,31,32^ and exert strong control over the timing of windows of developmental plasticity^3,14,15,33^. Previous work has shown that increases in the strength of glutamatergic thalamic inputs onto PV interneurons can enhance cortical plasticity^10^, likely by shifting the cortex’s excitation/inhibition balance to a plasticity-permissive state^3,4,10,14^. In contrast, the stabilization of thalamocortical-PV interactions by perineuronal nets^10^ or cell adhesion complexes^11^ serves to restrict ongoing plasticity^8^. Converging lines of evidence thus indicate that connectivity between the thalamus and cortex increases and then plateaus during postnatal cortical remodeling and maturation. This evidence implicates the strengthening and stabilization of thalamocortical connectivity in the opening and closing of biologically-programmed periods of cortical developmental plasticity.

While existing work identifies a role for thalamocortical axons in determining time windows of cortical malleability, this work was conducted almost exclusively in animal model sensory cortices that mature early in life. It is therefore not known whether coordinated maturation of cortical regions and thalamic structural pathways occurs in the human brain, with its evolutionarily expanded association cortices and uniquely protracted neurodevelopmental time course. This gap precludes a mechanistic understanding of whether age-related restructuring of thalamocortical connectivity could account for a defining feature of human neurodevelopment: the existence of S-A gradients of cortical plasticity. Here we assess whether maturational changes in thalamocortical structural connectivity progress from sensorimotor to association cortices and align with hierarchically-organized timescales of human cortical development. We first use diffusion MRI to create a new atlas of human thalamocortical connections, delineating over 200 structural connections between the thalamus and specific cortical regions. We then employ this atlas as an anatomical prior to identify consistent and regionally-specific thalamocortical connections in *N* = 2,676 youth across three independent discovery and replication datasets. Using this rich connectivity data, we test the hypothesis that windows of thalamocortical connection development unfold in time along the cortex’s S-A axis, with late maturation of association cortex connections coinciding with protracted refinement of higher-order cortices.

If thalamocortical connections are remodeled during childhood and adolescence^26,34,35^ in association with periods of experience-dependent cortical plasticity, the thalamus may also be a critical brain structure for determining windows of cortical environmental sensitivity. Indeed, work in animal systems has shown that a restructuring of thalamocortical inputs occurs in direct response to sensory environmental deprivation and precedes deprivation-induced cortical remodeling^12^. We therefore hypothesized that salient features of youths’ environments that affect cortical properties^2,36,37^ would furthermore show relationships with the strength of their thalamocortical connectivity— particularly for thalamic connections with regions in association cortex that have protracted developmental malleability. As described below, we demonstrate that the development of thalamocortical structural connectivity is spatiotemporally synchronized with environmentally-sensitive cortical developmental programs, thereby centering the thalamus in child and adolescent cortical development.

## Results

We evaluated a developmental model that relates the maturation of thalamocortical connectivity to cortical developmental variability and its environmental embedding across the S-A axis. To facilitate this evaluation, we leveraged one adult and three cross-sectional developmental datasets, advancements in diffusion MRI tractography, multi-modal maps of brain organization, and cortical charts of *in vivo* plasticity marker maturation. We first used high-resolution, multi-shell diffusion MRI data from the Human Connectome Project (HCP) Young Adult dataset (HCPYA; *N* = 1,065, ages 22-37 years) to create an atlas composed of > 200 white matter connections between the thalamus and individual cortical regions. We then applied this atlas to delineate person-specific thalamocortical connections in three independent developmental datasets. We used the Philadelphia Neurodevelopmental Cohort (PNC; *N* = 1,145, ages 8-23 years; a community-representative sample) and the Human Connectome Project in Development (HCPD; *N* = 572, ages 8-22 years; a typically developing sample) as primary discovery and replication datasets. The PNC and HCPD samples can be used to determine normative developmental trajectories in the age range when the S-A developmental hierarchy is most pronounced^2^. To characterize thalamocortical connectivity development in PNC and HCPD, we calculated fractional anisotropy (FA) in individual thalamocortical connections for all participants. We elected to study FA as it has been widely applied in the neurodevelopmental literature and can be appropriately derived from both single-shell (PNC) and multi-shell (HCPD) acquisitions. Finally, after obtaining our core set of results in PNC and HCPD, we examined the Healthy Brain Network (HBN; *N* = 959, ages 8-22 years; a psychiatric sample) to assess whether findings generalized to a help-seeking sample enriched for psychopathology. With these four datasets, we reproducibly delineate spatial gradients of thalamocortical structural connectivity features, temporal gradients of thalamocortical connectivity maturation, and associations between thalamocortical connectivity and the developmental environment.

### A human atlas of regionally-specific thalamocortical connections

Investigating whether the development of the cortex is constrained by the maturation of its structural connections with the thalamus necessitates the identification of white matter connections between the thalamus and localized cortical regions. However, reconstructing thalamocortical connections using diffusion MRI tractography is technically challenging. Obstacles include bottleneck effects that arise from the profusion of cortical fibers that enter and exit the thalamus, the need for thalamic connections to traverse areas of crossing fibers, the varied anatomy of thalamocortical connections, and typical reliance on tractography techniques that may overestimate cortical connectivity and sacrifice cortical endpoint specificity. We overcome these challenges by introducing a novel analytic approach executed in two steps: the creation of a new population-level atlas of human thalamocortical connections (step 1) that we subsequently leveraged as an anatomical prior to identify connections with the same trajectories in individuals’ data (step 2). When combined with a diffusion reconstruction method that resolves crossing fibers and parameter flexibility for tracking pathways with varied shapes, this atlas-based approach ultimately allowed for consistent identification of spatially-specific thalamocortical connections across individuals.

To create a high-resolution atlas of thalamocortical connections, we used a group-average diffusion template derived from 1,065 participants included in the HCPYA dataset. For this step, we chose to use the HCPYA diffusion data given that it was acquired in three shells with dense sampling and a high spatial resolution, making it well-suited to capture detailed thalamocortical connection anatomy. Moreover, we did not expect that entire thalamocortical connections would form or be pruned away between age 8 and young adulthood; rather, we anticipated changes solely in microstructural indices of connectivity strength. The HCPYA diffusion template was used to track between the thalamus and individual ipsilateral cortical regions defined by the HCP-multimodal parcellation (HCP-MMP). Following manual inspection and detailed curation of individual connections to ensure robustness and to remove spurious streamlines, we identified connections between the left and right thalamus and 238 individual cortical regions (119 bilaterally represented connections).

These connections covered 77% of the cortical mantle and spanned the entirety of the S-A axis (**Fig. 1a**; atlas connections colored by S-A axis rank). HCP-MMP regions where we did not identify thalamic connections were predominantly located in the cingulate, precuneus, insula, and lateral occipital cortex. Regions without thalamic connections had significantly smaller surface areas (anatomical enrichment test *p*_spin_ < 0.001; **Fig. 1b**) and significantly greater sulcal depth (anatomical enrichment test *p*_spin_ = 0.035; **Fig. 1c**); unconnected regions thus mainly occupied small cortical territories in sulcal banks.

**Figure 1.**
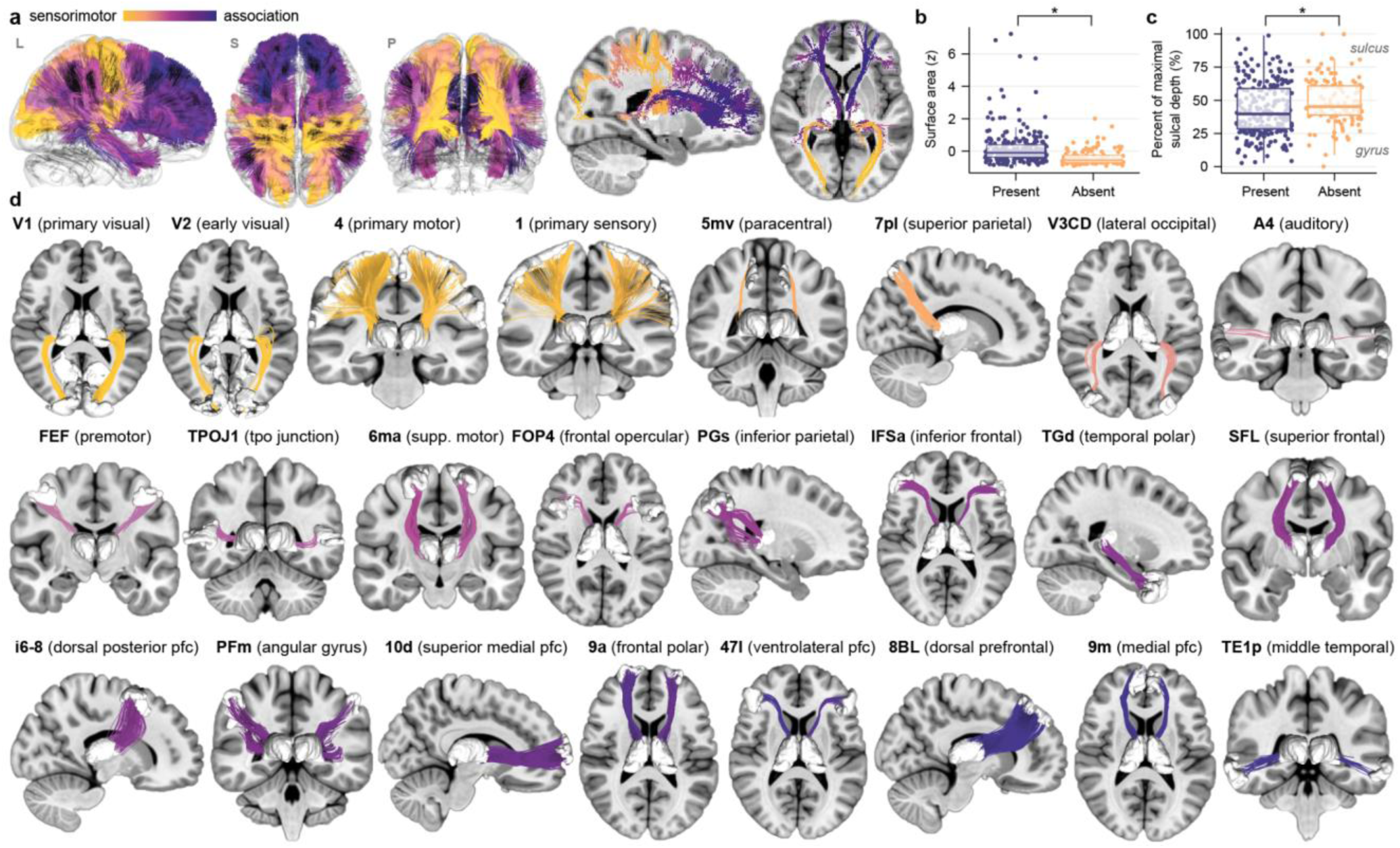
An atlas of regionally-specific thalamocortical structural connections. **a.** All connections that constitute the population-level thalamocortical connectivity atlas are shown in lateral (L), superior (S), and posterior (P) 3D views as well as 2D slice views. Connections are colored by the cortical endpoint’s position along the S-A axis of brain organization. The thalamocortical atlas exhibits extensive connectivity with cortical regions that occupy positions along the breadth of the S-A axis. **b.** Cortical regions that do not have thalamic connections represented in the atlas (absent) had significantly smaller surface area than included regions, as determined by a spatial permutation based null. Surface area z-scores are plotted for cortical regions with connections included (present) versus not included (absent) in the atlas. **c.** Cortical regions absent from the atlas had significantly greater sulcal depth, as determined by a null distribution generated using spatial permutations. Represented regions were more likely to be located in gyral crowns (0% of maximal sulcal depth) than in the banks of sulci (100% of maximal sulcal depth). In **c** and **d**, box plots summarizing data distributions are included (center line: median, hinges: first and third quartiles, whiskers: 1.5x interquartile range). **d.** Exemplar regionally-specific thalamocortical connections included in the atlas are shown. Connections are arranged from lowest (V1; yellow) to highest (TE1p; dark blue) position on the S-A axis.

The thalamocortical connections that make up the atlas have macroscale anatomical features that correspond to those identified in invasive tract tracing studies in macaques as well as non-invasive studies in humans that utilized advanced diffusion acquisitions^38^ and neuroanatomist-guided thalamic tracking pipelines^39^. For example, connections between the thalamus and primary visual cortex (V1 in **Fig. 1d**) show the Meyer’s loop of the optic radiation. Connections between the thalamus and primary somatosensory cortex (1 in **Fig. 1d**) exhibit fanning connectivity across the entirety of the medial-to-lateral somatosensory strip. Connections with supplementary motor regions (6ma and SFL in **Fig. 1d**) form crescent-shaped lamellae, in accordance with tract tracing studies^40^. Connections to the ventral lateral prefrontal cortex (IFSa and 47l in **Fig. 1d**) travel as compact stems along the anterior limb of the internal capsule and fan laterally to prefrontal cortex^38^. In sum, we developed a thalamocortical atlas of regionally-specific cortical connections that displays broad cortical coverage and anatomical accuracy. This atlas is made publicly available with implementation instructions for use in future studies (see Data Availability).

### Robust identification of thalamocortical connections in individuals

We used our thalamocortical connectivity atlas as a prior for an automated tractography approach to identify the same thalamocortical connections in PNC and HCPD participants’ data. This approach combines fiber tracking with trajectory-based connection recognition to accurately delineate atlas-defined thalamocortical structural connections in individual brains. As highlighted in **Fig. 2**, this approach consistently allowed for the identification of robust white matter pathways connecting the thalamus and specific cortical regions in participants of all ages in both PNC and HCPD. While the gross architecture of these connections was consistent across ages (**Fig. 2**), there were inter-individual differences in the strength of connectivity.

**Figure 2.**
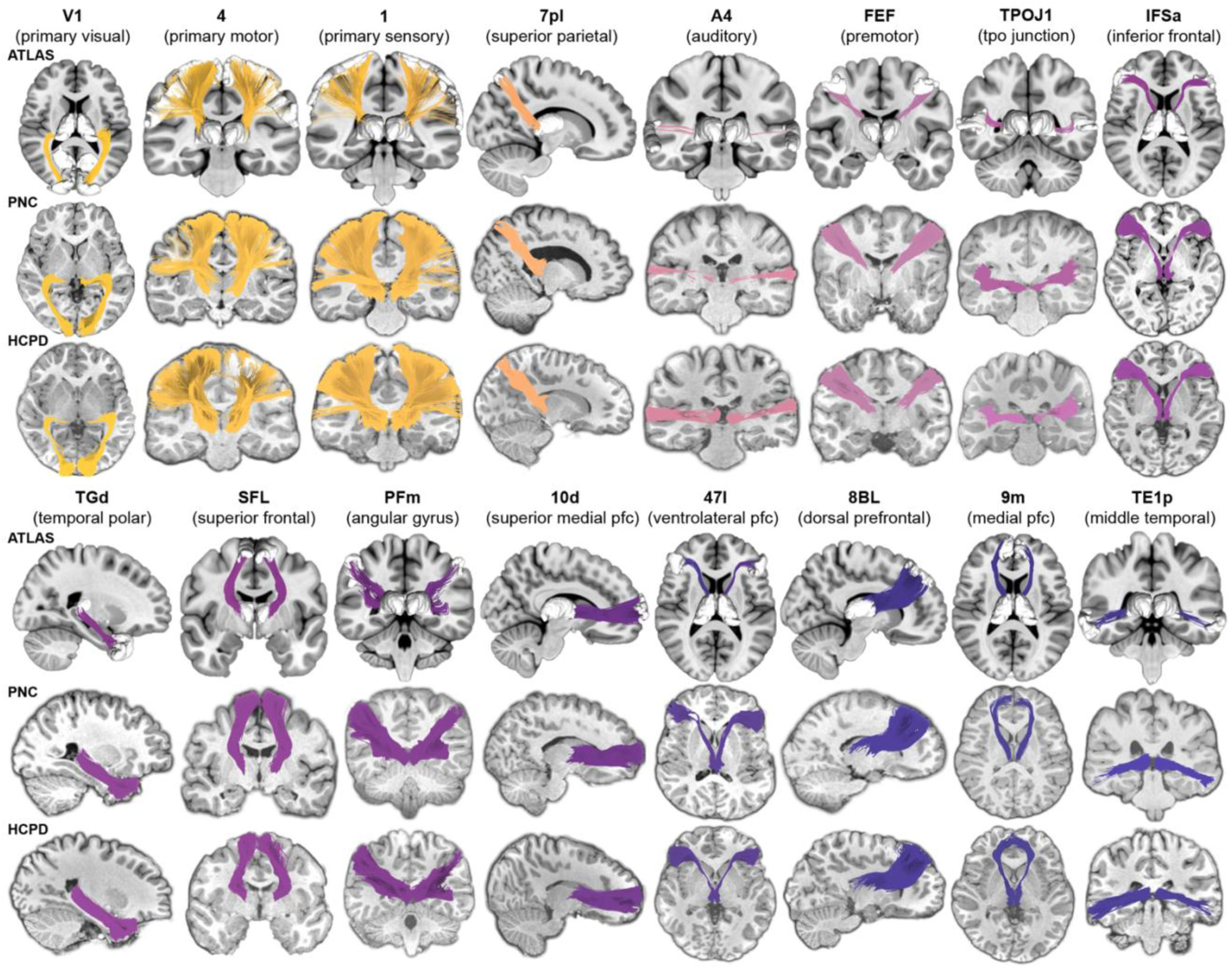
Thalamocortical structural connections are consistently reconstructed in individual participants. Exemplar thalamocortical connections included in the population-level atlas (*rows 1 and 4*) are shown here reconstructed in individual participants from PNC (*rows 2 and 5*) and HCPD (*rows 3 and 6*). Connection colors match those used in **Fig. 1** and reflect the S-A axis rank of the connected cortical region (yellow: lowest S-A ranks; purple/blue: highest S-A ranks). Person-specific connections showed remarkably high reconstruction robustness and anatomical endpoint accuracy. Each connection shown in the PNC and HCPD is from a different participant; a random number generator was used to select which participant’s data to show for each connection. The full distribution of ages is represented amongst the PNC data shown (minimum age in years = 8.3, 1^st^ quartile = 11.7, mean = 14.6, 3^rd^ quartile = 17.5, maximum age = 22.0) as well as the HCPD data shown (minimum age in years = 8.9, 1^st^ quartile = 11.0, mean = 14.9, 3^rd^ quartile = 18.6, maximum age = 21.8).

### Identified connections reflect thalamocortical circuit anatomy

Prior to using the connections reconstructed in PNC and HCPD participants to understand thalamocortical connectivity development, we aimed to further establish their anatomical validity. To accomplish this goal, we surveyed whether connections delineated in these datasets adhered to both the core-matrix structure of thalamic organization and the hierarchical arrangement of thalamocortical connection strength. Thalamic areas can be organized along a continuous core-matrix gradient based on the relative expression of more “core”-like or “matrix”-like neurons^30,41^. Core neurons are densest in first-order thalamic nuclei and send projections to sensory regions, whereas matrix neurons are prevalent in higher-order nuclei and connect to association regions. We therefore predicted that thalamocortical connections originating in more core versus matrix thalamus would project to different portions of the cortex’s S-A axis. We tested this differentiation by using a previously derived core-matrix thalamic gradient (C-M_t_; **Fig. 3a**)^30^ to assign each reconstructed connection a C-M_t_ value based on where its streamlines terminated within the thalamus. C-M_t_ values were calculated at the individual level and then averaged across participants in PNC and HCPD to derive group-level means.

**Figure 3.**
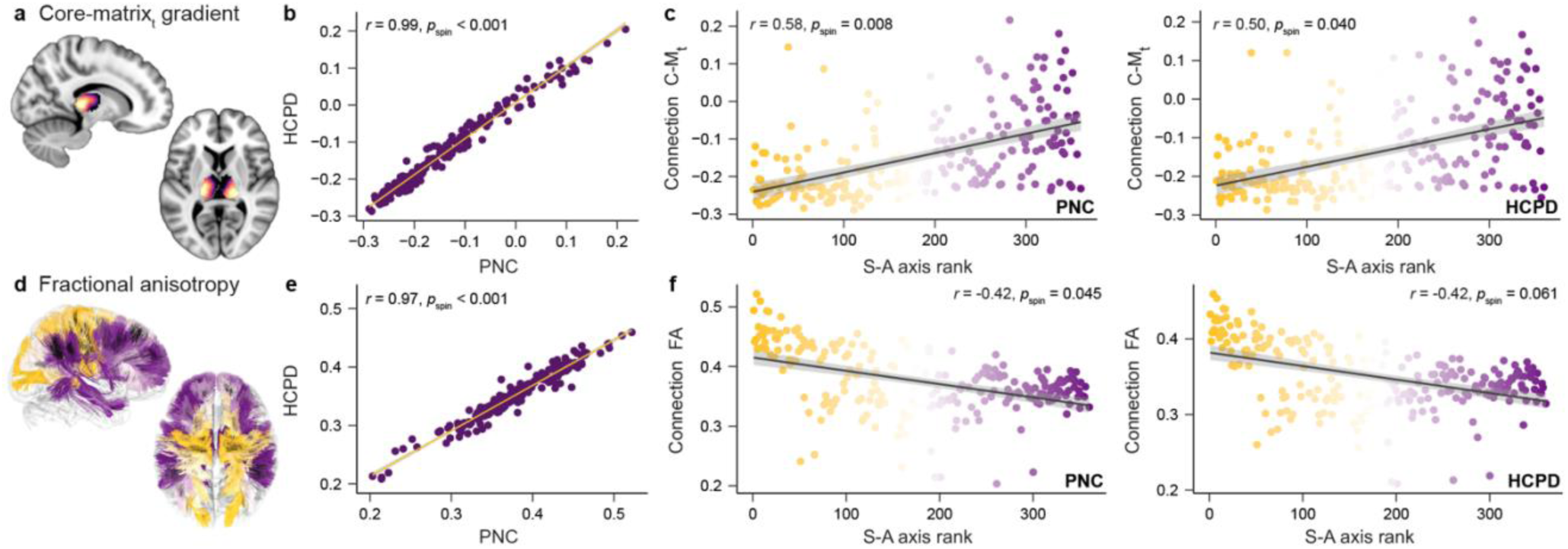
Identified structural connections reflect key features of thalamocortical circuit anatomy. **a.** The core-matrix thalamic (C-M_t_) gradient derived in prior work^30^ is shown in slices of the thalamus. Light yellow thalamic voxels were estimated to have the highest relative distribution of core neurons. Dark purple voxels were estimated to have the highest proportion of matrix neurons. **b.** We used the C-M_t_ gradient to assign each thalamocortical connection a value that indexes whether its streamlines terminated in thalamic areas with a higher percentage of core-like neurons (lower C-M_t_ values) or matrix-like neurons (higher C-M_t_ values). Connection-specific C-M_t_ values were nearly perfectly correlated between PNC and HCPD, serving as a general confirmation that delineated connections terminated in the same areas of the thalamus across datasets. **c.** Thalamic connection C-M_t_ values positively correlated with the sensorimotor-association (S-A) axis rank of the connection’s cortical partner in both PNC (*left*) and HCPD (*right*). Accordingly, both datasets showed evidence of core-to-sensory and matrix-to-association thalamocortical connectivity motifs. **d.** Thalamocortical connections are shown colored by mean FA (dark yellow: highest FA; dark purple: lowest FA; PNC data shown). **e.** Connection FA values were robustly correlated between PNC and HCPD, demonstrating that this structural connectivity feature is highly reproducible across youth samples. **f.** Thalamic connection FA values monotonically decreased along the S-A axis in PNC (*left*) and HCPD (*right*), revealing a continuum of connection strength and coherence that exhibits systematic hierarchical variation.

Connection-specific C-M_t_ values were nearly perfectly correlated between PNC and HCPD (*r* = 0.99, p_spin_ < 0.001; **Fig. 3b**), confirming that the atlas-constrained tractography approach generates reproducible profiles of thalamic connectivity. As predicted, the distribution of C-M_t_ values was not homogeneous across the S-A axis. Thalamic connections to the S-A axis’s sensorimotor pole originated in areas of the thalamus enriched with core neurons (lowest C-M_t_ values). Connections that originated in matrix-like thalamic areas (higher C-M_t_ values) were linked to the axis’s association pole. A distribution of increasing C-M_t_ values across the S-A axis was observed in both PNC (*r* = 0.58, *p*_spin_ = 0.008) and HCPD (*r* = 0.50, *p*_spin_ = 0.040) and provides evidence that reconstructed thalamocortical connections exhibit well-described core-sensory and matrix-association connectivity motifs (**Fig. 3c**).

Thalamic pathways that project to sensorimotor versus association cortices are also known to differ in their microstructural anatomy. Thalamic connections to primary cortex are dense, strong, and project in a spatially constrained manner, whereas projections to association cortex are sparser and more spatially diffuse^30,41^. We therefore predicted that FA, a microstructural measure that increases with connection density and coherence, would be highest for thalamic connections with primary sensory regions and decrease along the S-A axis. As for C-M_t_ values, we calculated FA for every thalamocortical connection at the individual participant level and computed a group-level connection mean for PNC and HCPD (**Fig. 3d**). Connection-specific FA values were highly reproducible between PNC and HCPD (*r* = 0.97, p_spin_ < 0.001; **Fig. 3e**). In line with our prediction, connection-specific FA values negatively correlated with the S-A axis rank of the connection’s cortical partner (PNC: *r* = -0.42, *p*_spin_ = 0.045; HCPD: *r* = -0.42, *p*_spin_ = 0.061; **Fig. 3f**), suggesting graded changes in connection microstructure along this organizational axis. The current set of findings confirms that reconstructed pathways intrinsically reflect established thalamic cellular classifications and cortical connection profiles.

### Thalamocortical connections show a spectrum of developmental change

Having now demonstrated that our atlas-based approach extracts connections with properties that capture thalamocortical circuit anatomy, we sought to investigate whether these connections exhibit hierarchically-organized variability in developmental timing. We began by using generalized additive models (GAMs; accounting for sex and head motion) to characterize age-dependent trajectories of FA for all connections. FA significantly increased in the majority of thalamocortical connections during childhood and adolescence, with 90% (PNC) and 78% (HCPD) of connections showing significant (*p*_FDR_ < 0.05) developmental effects. Although most connections showed a general increase in FA, a spectrum of developmental trajectories could be seen in both datasets (Fig. 4a), paralleling modes of developmental variability that typify the cortex^1^. As a result of these variable trajectories, the magnitude of GAM-derived age effects (quantified as the partial *R^2^*) differed across connections (Fig. 4b). Age effects were strongly correlated between PNC and HCPD (*r* = 0.73 *p*_spin_ < 0.001; Fig. 4c).

**Figure 4.**
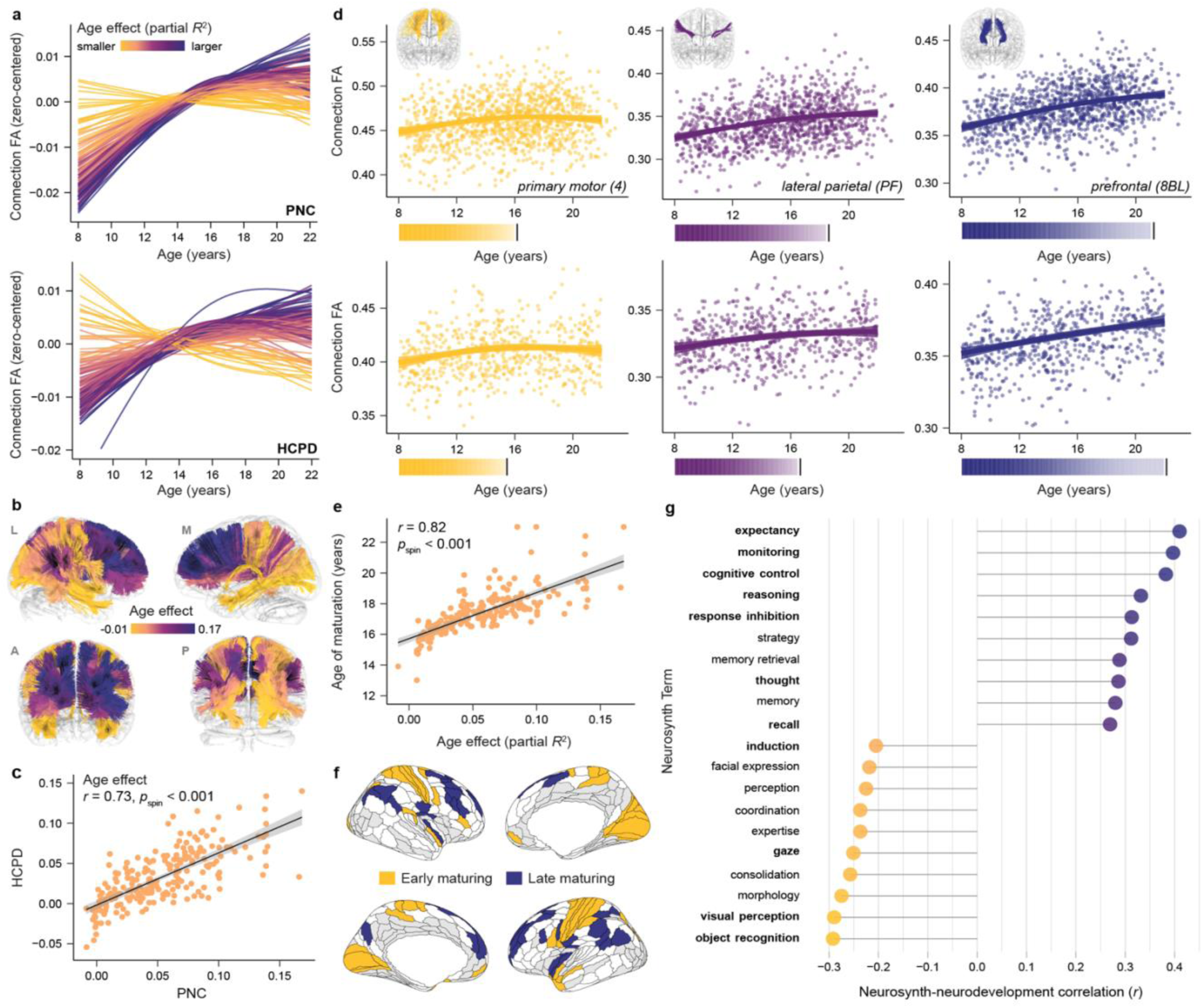
Charting variability in the magnitude and timing of thalamocortical structural connectivity development. **a.** Fractional anisotropy (FA) developmental trajectories (zero-centered GAM smooth functions) are displayed for right hemisphere thalamocortical connections in PNC (*top*) and HCPD (*bottom*). Connections are colored by their statistical age effect (partial *R^2^*). A clear neurodevelopmental spectrum is observable in both datasets. **b.** Thalamocortical connections from the atlas are colored by their age effect (PNC data), revealing the brain-wide distribution of developmental heterogeneity. **c.** A correlation plot confirming close correspondence between connection-specific age effects derived in PNC and HCPD. **d.** Connection-specific FA trajectories that exemplify differences in the magnitude and timeframe of developmental change are shown, overlaid on participant-level data for PNC (*top*) and HCPD (*bottom*). Developmental trajectories represent GAM-predicted FA values with a 95% credible interval band. The corresponding color bars chart the rate of increase in FA during windows of significant developmental change and demarcate ages of connection maturation. **e.** GAM-derived age effects and ages of maturation were correlated in both datasets (PNC shown), revealing that the age of maturation metric provides insight into both the extent and timing of development. **f.** A brain map localizing cortical regions with the earliest maturing thalamic connections (age of maturation first quartile; yellow) and latest-maturing thalamic connections (fourth quartile; blue) is shown. White designates cortical regions with connections to the thalamus that matured in middle age quartiles. Light grey indicates regions with connections not included in the atlas. **g.** Results of an analysis that correlated the map of thalamocortical maturational age with psychological term maps from Neurosynth. Psychological terms associated with cortical regions that have thalamic connections that mature at the youngest ages (negatively correlated terms; yellow) and oldest ages (positively correlated terms; blue) are shown. PNC data is presented. Terms that were additionally included in the list of the 10 most negatively or 10 most positively correlated terms in HCPD are bolded.

Connections between the thalamus and sensory or motor regions tended to show the smallest age effects and to exhibit the shortest windows of developmental change, as seen for the connection to primary motor area 4 (Fig. 4d**, yellow**). Thalamic connections with multimodal association cortices (e.g., area PF of the supramarginal gyrus; Fig. 4d**, purple**) tended to develop for relatively longer. Thalamocortical connections exhibiting the largest and most protracted developmental increases involved regions of the lateral prefrontal and parietal association cortex, as exemplified by the thalamic connection to superior prefrontal region 8BL (Fig. 4d**, blue**). We assessed whether these observed developmental trajectories differed by sex. Although a small subset of connections showed potentially diverging trajectories between males and females starting in the mid-teens, nearly all age-by-sex interactions were not significant (0% of connections significant in PNC and < 2% significant in HCPD). Accordingly, subsequent results model a single trajectory across sexes.

To quantitatively study differences in developmental timing, we computed the age at which each thalamocortical connection matured. Maturational age was operationalized as the age at which developmental change in FA (the first derivative of the age spline) was no longer significantly different from 0, denoting a plateau in the developmental trajectory. Connection-specific ages of maturation were highly similar between the two datasets (*r* = 0.57, *p*_spin_ < 0.001) and furthermore correlated with connection-specific age effects (partial *R^2^*) within each dataset (Fig. 4e). Notably, large differences in maturational timing emerged across thalamocortical connections in both datasets: the relative difference in maturational age between the earliest and latest maturing connections was 9.9 years in PNC and 11.1 years in HCPD. Identifying thalamic connections that matured at the youngest ages (first quartile; Fig. 4f, yellow) versus the oldest ages (fourth quartile; Fig. 4f, blue) differentiated primary and early visual, somatomotor, and auditory regions from lateral prefrontal and parietal regions.

We next sought to put these observed differences in connection maturational timing in a behavioral context. To do so, we used prior task-based fMRI results—amassed and meta-analyzed via Neurosynth—to identify the psychological functions subserved by cortical regions with thalamic connections that matured at younger versus older ages. We first mapped the maturational ages of all connections to the cortex. We then computed the correlation between this maturational map and psychological term meta-analytic maps for 123 terms included in the Cognitive Atlas (producing 123 independent correlations between neurodevelopment and Neurosynth variables). In this analysis, a negative correlation between the thalamocortical maturational map and a psychological term map indicates that the psychological term is linked to regions with early maturing thalamic connections. In contrast, positive correlations identify psychological functions that can be ascribed to regions with late maturing connections. We thus identified the 10 most negatively and positively correlated terms in both PNC and HCPD and found that 11 of these 20 developmentally-relevant terms overlapped between datasets (bolded terms in Fig. 4g; p_PERM_ < 0.001 in a term-overlap permutation analysis). Psychological terms linked to cortical regions with early-maturing thalamic connections predominantly described sensory and motor processing and object classification functions (e.g., visual perception, coordination, object recognition; Fig. 4g negative correlations). Cognitive terms linked to cortical regions with late-maturing thalamic connections evoked executive control, decision-making, and information retrieval functions (e.g., cognitive control, reasoning, recall; Fig. 4g positive correlations). Overall, these results establish that thalamocortical structural connections exhibit different timescales of development. Maturational timing diverges most between connections to sensorimotor cortices that execute externally oriented functions and those to association cortices that are essential for higher-order cognitive control.

### Thalamocortical connection maturation unfolds along the S-A axis

If connections between the thalamus and the cortex play a role in organizing differences in developmental timing across the S-A axis, we would expect observed variability in thalamocortical connection maturation to systematically align with this axis. To study this alignment, we demarcated age windows of significant developmental change for each connection and visualized whether the length of these windows increased between the S-A axis’s sensorimotor and association poles. As shown in Fig. 5a, connection-specific windows of developmental change were staggered in time across the S-A axis and were most protracted for thalamic connections with transmodal association regions. To test whether this developmental pattern emerged due to connections maturing at progressively older ages along the S-A axis, we calculated the correlation between each connection’s age of maturation and its S-A axis rank (Fig. 5b, c). Lending strong support to our primary developmental hypothesis, we found that the age of connection maturation progressively increased for connections to cortices ranked higher in the S-A axis. This positive correlation between S-A axis ranks and connection-specific ages of maturation was similar in strength in PNC (*r* = 0.49, *p*_spin-FDR_ = 0.004) and HCPD (*r* = 0.51, *p*_spin-FDR_ = 0.008), underscoring that this spatiotemporal developmental pattern unfolds in independent samples (Fig. 5d).

**Figure 5.**
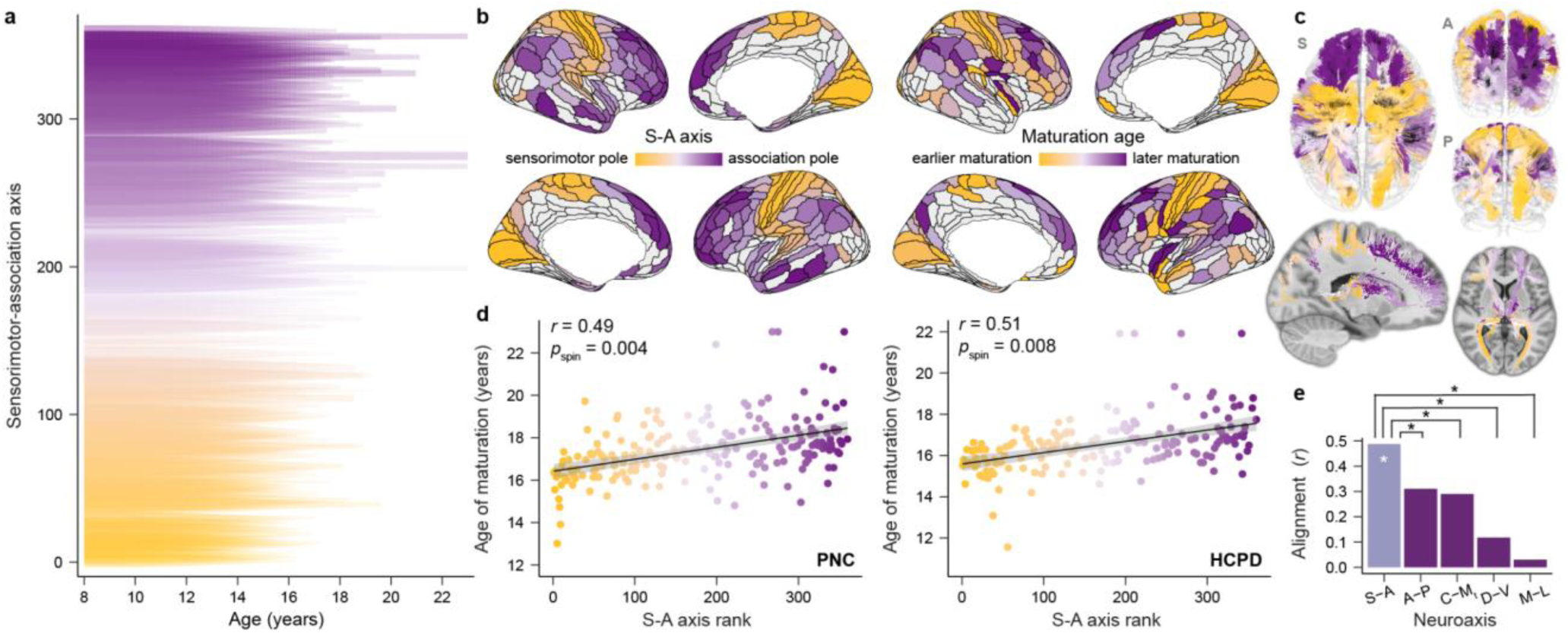
Thalamocortical structural connections mature at progressively older ages along the S-A axis. **a.** Age windows during which thalamocortical connections showed significant increases in FA are shown for every individual connection (PNC data). Connection-specific developmental windows are ordered along the y-axis and colored by the sensorimotor-association (S-A) axis rank of the connection’s cortical endpoint. Periods of significant developmental change were derived from the first derivative of each connection’s GAM smooth function for age, which quantifies the rate of change in FA at a given age. Significant derivative values (as determined by their simultaneous 95% confidence interval) are plotted here to delineate developmental windows; the relative transparency and linewidth of the derivative reflects the age-specific magnitude of developmental change. **b.** The maturational map depicting the age at which each cortical region’s thalamic connection matured (*right*) exhibits shared spatial topography with the S-A axis (*left*). Light grey regions in these cortical maps were not represented in the thalamocortical atlas. **c.** Thalamocortical connections from the tractography atlas are colored by the connection’s age of maturation to further illustrate the spatial structure of developmental effects. **d.** Ages of thalamic connection maturation systematically vary along the S-A axis in both PNC (*left*) and HCPD (*right*). Thalamocortical connections to the axis’s association pole tended to mature latest. **e.** Results of an analysis comparing the alignment of thalamocortical connectivity maturational timing to the S-A axis versus major cortical and thalamic axes. This analysis revealed greater alignment to the S-A axis than to anterior-posterior (A-P), dorsal-ventral (D-V), and medial-lateral (M-L) cortical axes as well as the core-matrix thalamic (C-M_t_) gradient (PNC data shown). Whereas the correlation between the maturational map and the S-A axis was significant (white star; data presented in panel **d**), spatial permutation tests confirmed that correlations between the maturational map and A-P, D-V, and M-L axes and the C-M_t_ gradient were not significant. Statistical comparisons of correlations further demonstrated that the correlation with the S-A axis was significantly greater in magnitude than correlations with these four neuroaxes (black stars).

To evaluate the specificity of these findings, we tested whether across-connection differences in ages of maturation were best captured by the S-A axis, or could be equally or better explained by other large-scale cortical or thalamic axes. Specifically, we assessed how connectivity maturational timing varied along anterior-posterior (A-P), dorsal-ventral (D-V), and medial-lateral (M-L) cortical axes and the core-matrix thalamic gradient (Fig. 5e). Correlations between the age of thalamocortical connection maturation and A-P (*r* = 0.31), D-V (*r* = 0.12), and M-L (*r* = 0.03) cortical axes and the C-M_t_ gradient (*r* = 0.29) were not significant in the PNC (all *p*_spin-FDR_ > 0.05). Furthermore, statistical tests for comparing the magnitude of two dependent, overlapping correlations indicated that connection maturational ages were significantly more correlated with the S-A axis than with A-P, D-V, and M-L axes and the C-M_t_ gradient (*p*_FDR_ < 0.001 for all four correlation comparisons). We observed the same results in HCPD, with strongest alignment to the S-A axis as compared to all other cortical and thalamic neuroaxes (correlation comparison for A-P: *p*_FDR_ = 0.124; D-V: *p*_FDR_ < 0.001; M-L: *p*_FDR_ < 0.001; C-M_t_: *p*_FDR_ = 0.029).

### Coordinated development of thalamocortical connections and cortical plasticity

The above results reinforce past findings that the S-A axis can be understood as a principal axis of child and adolescent neurodevelopment and indirectly relate thalamocortical connectivity maturation to temporal variation in cortical development. We therefore next endeavored to directly assess correspondence between the maturation of thalamocortical structural connections and three non-invasive and biologically linked readouts of cortical developmental plasticity. Animal studies have shown that the maturation of PV inhibitory interneurons^14,33^ and the formation of intracortical myelin^42^ serve as biological regulators of critical periods of plasticity. As interneurons strengthen their outputs and myelin is formed, there is a reduction in the cortex’s excitation/inhibition (E/I) ratio and a consequent suppression and sparsification of intrinsic cortical activity^2,43,44^. The transition from malleable to mature cortex can therefore be indexed by three signatures of decreasing circuit plasticity: a decline in the E/I ratio, an increase in cortical myelin content, and a reduction in the amplitude of intrinsic cortical activity. We explored whether the timing of thalamocortical connection maturation was temporally coordinated with the sensorimotor-to-associative development of these three readouts of shifting cortical plasticity.

We used developmental maps charting how *in vivo* measures sensitive to the cortical E/I ratio^5^, cortical myelin content^6^, and cortical intrinsic activity amplitude^2^ change with age during childhood and adolescence. We discovered that cortical regions with thalamic connections that developed for longer also exhibited smaller developmental declines in the E/I ratio (Fig. 6a), experienced a slower rate of intracortical myelin growth (Fig. 6b), and showed an initial decrease in intrinsic fluctuation amplitude at older ages (Fig. 6c). As such, protracted maturation of thalamocortical connections was associated with extended expression of neurochemical, structural, and functional markers indicative of higher circuit plasticity. Correlations between dataset-specific thalamocortical connectivity maturation maps and non-invasively estimated developmental maps of E/I ratio, cortical myelin, and intrinsic activity were significant in all cases in both datasets: PNC (model-derived E/I ratio: *r* = 0.45, *p*_spin-FDR_ < 0.001; T1/T2 ratio: *r* = -0.45, *p*_spin-FDR_ = 0.001; BOLD fluctuation amplitude: *r* = 0.30, *p*_spin-FDR_ = 0.030) and HCPD (model-derived E/I ratio: *r* = 0.45, *p*_spin-FDR_ < 0.001; T1/T2 ratio: *r* = -0.43, *p*_spin-FDR_ = 0.003; BOLD fluctuation amplitude: *r* = 0.41, *p*_spin-FDR_ = 0.009) (Fig. 6). These relationships provide evidence that the development of thalamocortical structural connectivity and cortical plasticity is spatiotemporally tethered.

**Figure 6.**
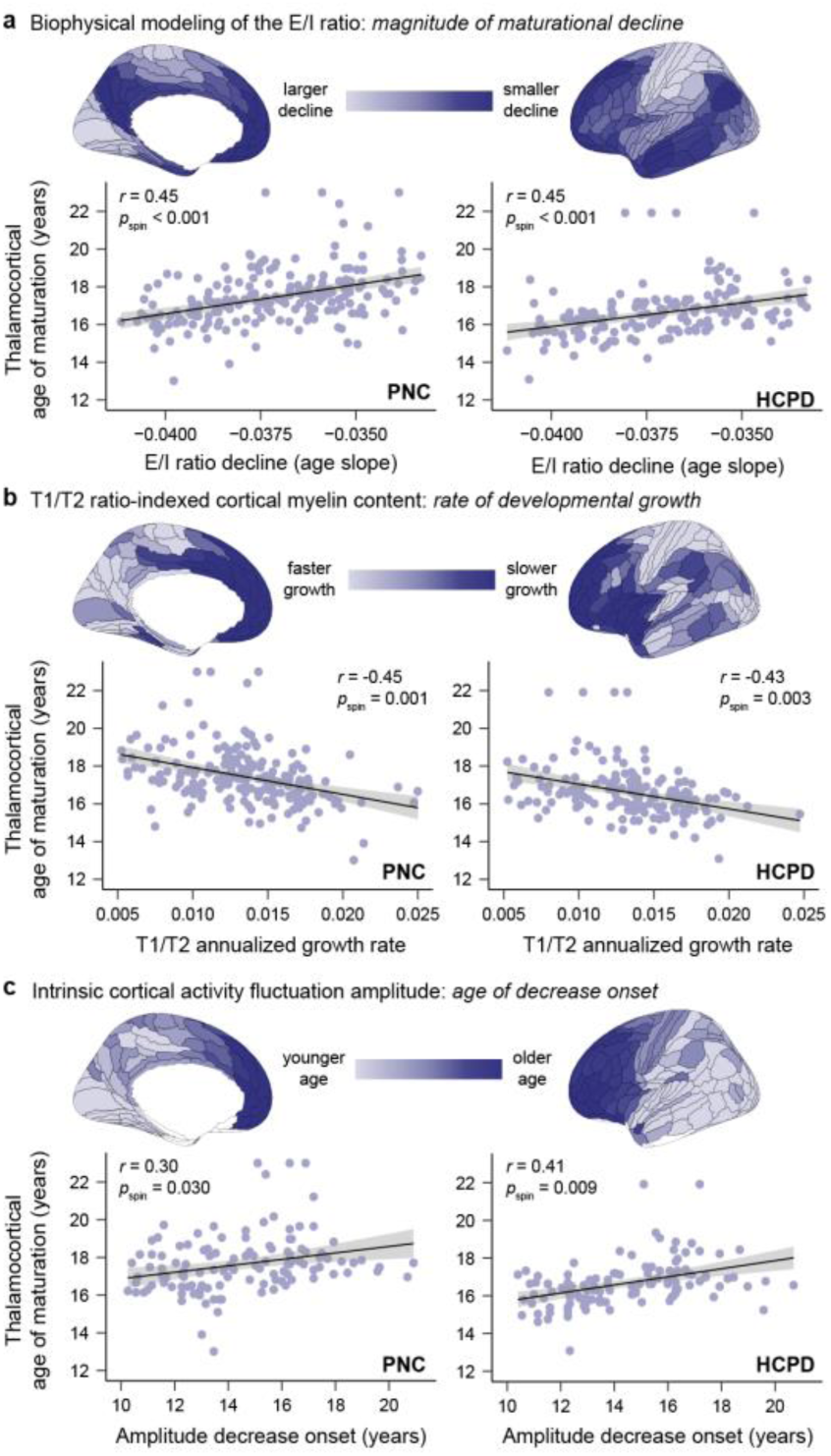
Thalamocortical structural connection maturation synchronizes with timescales of cortical plasticity. Maps of thalamocortical connection maturation computed from PNC (*left column*) and HCPD (*right column*) diffusion data correlate with brain charts of child and adolescent cortical development. Cortical maps charting the development of E/I ratio (**a**), cortical myelin (**b**), and intrinsic activity amplitude (**c**) are shown. In all three cortical maps, darkest blue brain regions are those that express signatures of protracted developmental plasticity. **a**. Cortical regions with thalamic connections that mature at older ages undergo smaller age-related reductions in the E/I ratio during childhood and adolescence (less negative age slopes), implying that they remain in a relatively less mature, plasticity-permissive state for longer. E/I ratio was estimated in developmental data in Zhang, Larsen, et al.^5^ by applying a biophysically plausible circuit model to resting state functional MRI data. **b**. Cortical regions with thalamic connections that mature at older ages show a slower annualized growth rate of T1/T2 ratio-indexed cortical myelin content, suggesting that they experience slower maturation of a structural feature that restricts developmental plasticity. T1/T2 ratio development data is from Baum et al., 2022^4^. **c**. Cortical regions with thalamic connections that mature at older ages exhibit later-onset declines in the amplitude of intrinsic activity fluctuations, indicative of temporally delayed reductions in a putative functional signature of developmental plasticity. The age at which intrinsic activity amplitude began to decrease in each cortical region was determined by Sydnor et al., 2023^2^ through developmental modeling of age-related changes in BOLD fluctuation amplitude.

### Environmental sculpting of thalamocortical connectivity across the S-A axis

Ample prior work has documented strong associations between socioeconomic features of the environment and cortical properties during youth. During infancy^36^, childhood^45^, and adolescence^2^, across-cortex variation in environment associations has been found to be systematically patterned along the S-A axis. In a series of analyses, we aimed to ascertain whether a similar principle governs interactions between youths’ environmental exposures and thalamocortical structural connectivity. Specifically, we studied relationships between household and neighborhood indicators of socioeconomic conditions and thalamocortical connection FA across the S-A axis. Household socioeconomic status was proxied by caregiver years of education in PNC and by both caregiver education and income-to-needs ratio in HCPD. Neighborhood-level socioeconomic information was only available in the PNC and was summarized via factor analysis of geocoded census data. The factor analysis generated person-specific neighborhood factor scores; a higher factor score indicates that a child lived in a neighborhood with a higher percentage of residents who were married, employed, and high-school educated and a lower percentage of residents in poverty.

We first modeled associations between indicators of socioeconomic position and thalamocortical connection FA using GAMs that accounted for developmental effects. Over half (57%) of thalamocortical connections showed significant relationships between connection FA and neighborhood environment factor scores (*p*_FDR_ < 0.05) in the PNC. In contrast to these robust effects, only 7% of connections showed a significant association (*p*_FDR_ < 0.05) with caregiver education in PNC. When caregiver education and neighborhood factor scores were entered into the same model as part of a specificity analysis, all significant caregiver education effects were abolished. Conversely, 91% of connections showing a significant neighborhood-level effect still exhibited a significant association with connection FA. Mirroring these null household-level findings in PNC, no thalamocortical connections showed a significant relationship between FA and either caregiver education or income-to-needs ratio in HCPD (all *p*_FDR_ > 0.05).

In the PNC, significant relationships between connection FA and the neighborhood environment factor score were widely distributed across thalamocortical connections and across the S-A axis (Fig. 7a). Environment effects (*t* values) were nearly exclusively positive (93% positive), indicating that more socioeconomically advantaged neighborhoods were associated with higher connection FA. Environment factor scores were not correlated with diffusion scan mean framewise displacement (*r* = 0.04, *p* = 0.202), supporting that these associations were not driven by motion. To better understand the nature of these FA-environment associations, we modeled the maturation of thalamocortical connection FA for low and high factor scores for five quintiles of the S-A axis. These environmentally-stratified developmental trajectories showed that lower neighborhood factor scores were associated with lower FA throughout the course of child and adolescent development in all portions of the S-A axis (Fig. 7b).

**Figure 7.**
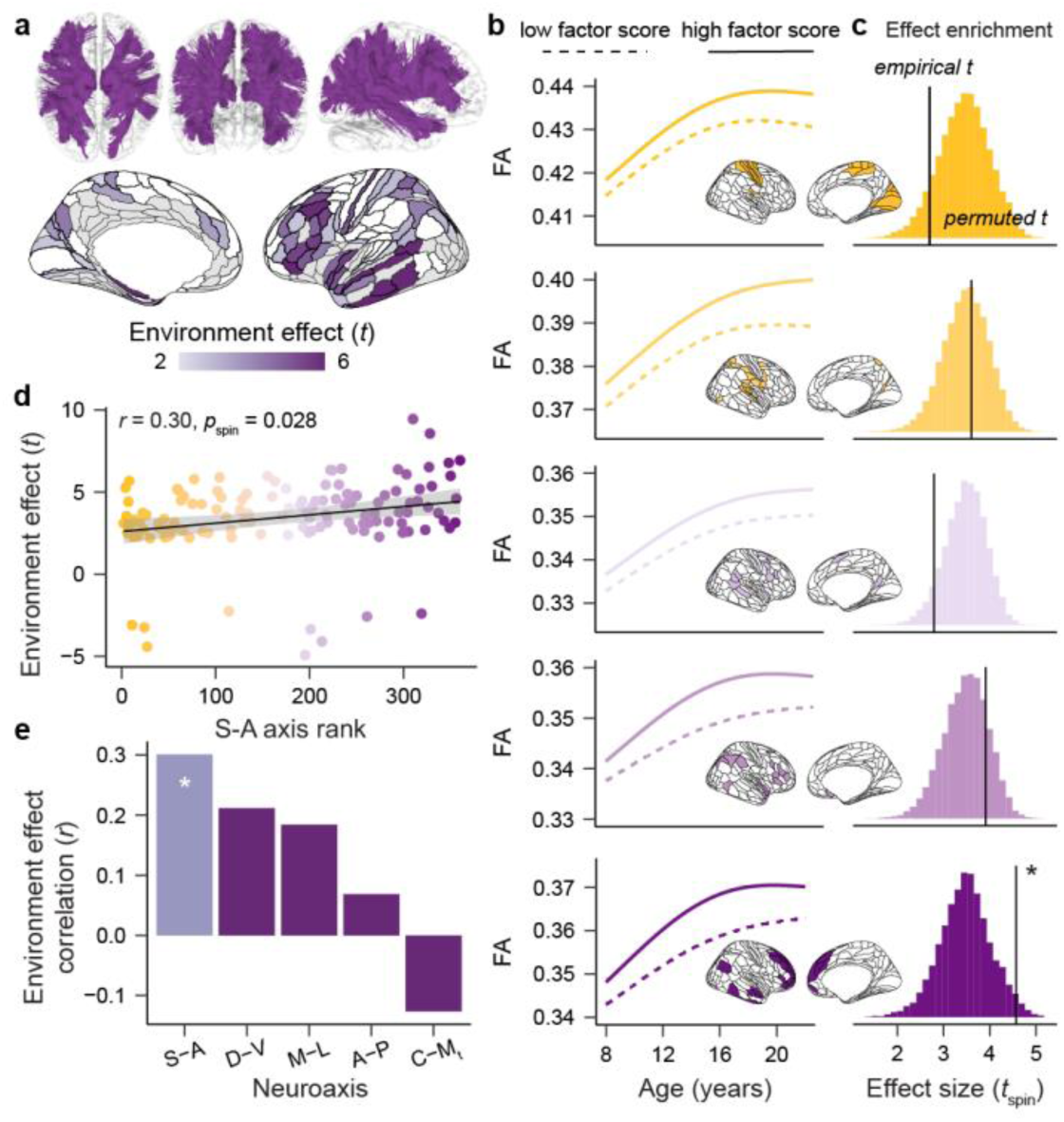
Hierarchically-organized relationships between the neighborhood environment and thalamocortical structural connectivity. **a.** Thalamocortical connections that showed a significant association between neighborhood environment factors scores and connection FA are shown in purple (*top*). The statistical effect (*t* value) associated with each significant connection is displayed on the cortical surface (*bottom*). Significant effects were present across much of the cortex and were strongest in lateral frontal and temporal cortices. White and grey cortical regions denote regions with connections that had non-significant environment effects or that were not analyzed, respectively. **b.** GAM-predicted trajectories of FA development are displayed for low (10^th^ percentile) and high (90^th^ percentile) factor scores for thalamic connections to five quintiles of the sensorimotor-association (S-A) axis. Trajectories model environment-related differences in connection FA from childhood to early adulthood. **c.** Results of the environment effect enrichment analysis are displayed for five quintiles of the S-A axis. This analysis uncovered that neighborhood environment *t* values were significantly greater in magnitude for thalamic connections to the fifth quintile of the S-A axis (darkest purple; *bottom*) when compared to connections with the rest of the cortex. For each of the five quintiles of the S-A axis, the empirical mean *t* value in that quintile is indicated with the black line along with the null distribution of permuted mean *t* values (*t*_spin_) obtained from 10,000 spatial permutations of the cortex-projected environment effect map. **d.** Thalamic connections to cortical regions ranked higher in the S-A axis showed relatively larger associations between connection FA and neighborhood environment factor scores, resulting in a significant correlation between S-A axis ranks and thalamocortical environment effects. A small number of negative *t* values were observed in motor, premotor, and orbitofrontal cortex. **e.** Results of an analysis correlating each connection’s environment effect (from **a**, *bottom*) with its position in the S-A axis as well as major cortical axes and the core-matrix thalamic gradient. Only the correlation with the S-A axis was significant (white star; *p*_spin-FDR_ < 0.05).

Although neighborhood environment associations were present across the S-A axis, the magnitude of significant effects was not homogeneous across connections: larger effects could be seen in thalamic connections to lateral prefrontal and lateral temporal cortices (Fig. 7a, bottom). We therefore conducted an analysis to test whether environment effects (*t* values) were statistically enriched for connections to the association end of the S-A axis. Enrichment tests for five quintiles of the S-A axis uncovered that the strongest neighborhood environment effects (high *t* values) were indeed overrepresented for thalamic connections with the association pole of the S-A axis (fifth quintile enrichment analysis: *p*_spin_ = 0.025; Fig. 7c). Substantiating this finding of relatively larger effects at the association pole, a second analysis correlating regional S-A axis ranks with the cortex-projected map of significant neighborhood environment effects confirmed a significant, positive association (*r* = 0.30, *p*_spin-FDR_ = 0.028; Fig. 7d). Specificity analyses revealed that alignment was significantly stronger to the S-A axis than to the A-P axis (correlation comparison *p*_FDR_ = 0.001) and the C-M_t_ thalamic gradient (correlation comparison *p*_FDR_ < 0.001) and that correlations between environment effect statistics and A-P, M-L, and D-V cortical axes and the C-M_t_ gradient were all non-significant (all *p*_spin-FDR_ > 0.05) (Fig. 7e). These analyses demonstrate that neighborhood-level socioeconomic conditions relate to thalamocortical connectivity properties during youth, with connections that experience protracted development displaying the greatest environmental sensitivity.

### Results generalize to a youth sample enriched for psychopathology

Thus far, we have demonstrated that developmental and environmental influences on thalamocortical connection properties vary depending on a connection’s position in the cortex’s S-A axis. In a final set of analyses, we investigated whether these findings generalize to a clinical sample recruited with the goal of representing transdiagnostic youth psychopathology. The HCPD sample was designed to study “typical” brain development. The PNC used community-based recruitment and was not specifically enriched for psychopathology. In contrast, the Healthy Brain Network is a study of help-seeking youth where approximately 85% meet criteria for a clinical diagnosis.

In HBN, 74% of thalamocortical structural connections showed a significant developmental change in FA (*p*_FDR_ < 0.05), with developmental profiles substantially varying across connections (Fig. 8a). Connection-specific age effects obtained in HBN strongly and significantly correlated with those obtained from the PNC (*r* = 0.73, *p*_spin_ < 0.001; Fig. 8b), further underscoring that our results capture a generalizable developmental signature. Neurosynth-based decoding of connection maturational timing linked early-maturing thalamocortical connections to perceptual and motor functions and late-maturing connections to memory retrieval, decision making, and cognitive control (Fig. 8c; 11 overlapping terms with PNC). Thalamocortical connections exhibited a hierarchical maturational gradient. As a result, connection maturational age was correlated with the S-A axis (*r* = 0.69, *p*_spin_ = 0.002; Fig. 8d). Furthermore, ages of connection maturation aligned with age-related change in the three neuroimaging-based readouts of cortical developmental plasticity (model-derived E/I ratio: *r* = 0.57, *p*_spin-FDR_ = 0.007; T1/T2 ratio: *r* = -0.58, *p*_spin-FDR_ = 0.015; BOLD fluctuation amplitude: *r* = 0.69, *p*_spin-FDR_ = 0.007; Fig. 8e).

**Figure 8.**
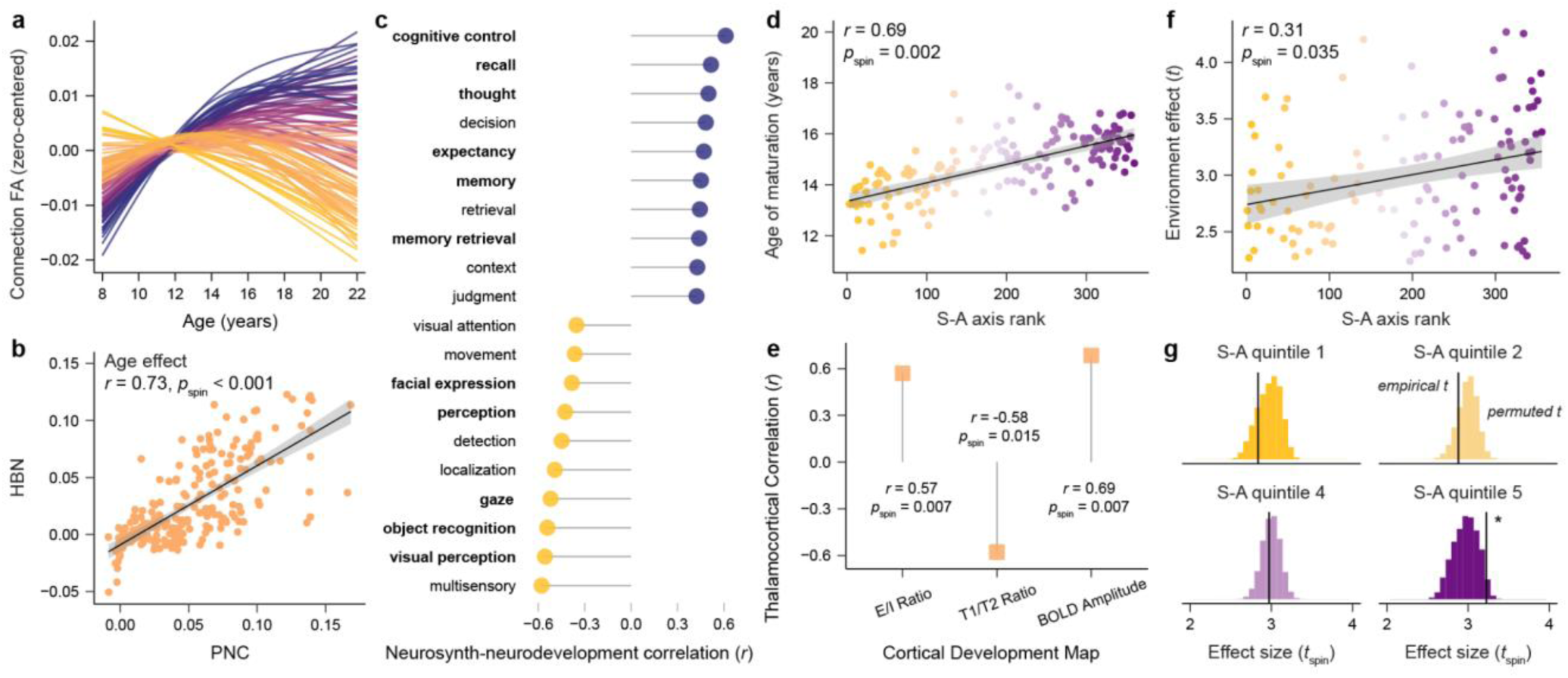
Developmental and environmental results are generalizable to youth with psychopathology. An overview of key results from the HBN sample, a clinical sample of youth that is enriched for psychopathology. **a.** Structural connections between the thalamus and cortex exhibit heterogenous profiles of fractional anisotropy (FA) development. **b.** Connection-specific age effects derived in HBN correlate with those obtained in the PNC. **c.** Neurosynth-based contextualization of thalamocortical connection developmental timing reveals psychological functions associated with cortical regions with early-maturing thalamic connections (negatively correlated terms) and late-maturing thalamic connections (positively correlated terms). Bolded terms overlap with those identified in PNC. **d.** The age at which thalamocortical connections mature progressively increased for connections to cortical regions located higher in the S-A axis, resulting in a positive correlation between ages of thalamic pathway maturation and S-A axis ranks. **e.** Thalamocortical connection maturation significantly correlated with non-invasively derived maps charting the development of cortical properties, including the development of the cortical excitation/inhibition (E/I) ratio, cortical T1/T2 ratio, and cortical BOLD activity fluctuation amplitude. The strength and significance of each of these three correlations is indicated. **f.** A plot depicting the spread of significant neighborhood environment effects (statistical *t* values) across the S-A axis is shown. Positive environment effects indicate that more socioeconomically advantaged neighborhood environments were associated with higher thalamocortical connection FA. Significant environment associations were found across the entire S-A axis. Effects became significantly larger when moving towards the axis’s association pole. **g.** The environment enrichment analysis confirmed that neighborhood environment effects were significantly greater in magnitude for thalamic connections to the fifth quintile of the S-A axis as compared to connections with the rest of the cortex.

We used the same geocoding-based factor analysis approach as in the PNC to summarize information about each participant’s neighborhood-level socioeconomic circumstances. In HBN, 53% of thalamocortical connections exhibited a significant relationship between neighborhood environment factor scores and connection FA. All associations were positive, linking more advantaged neighborhoods (higher factor scores) to stronger thalamocortical connectivity. As in the PNC, the magnitude of environment-connectivity associations increased in strength along the S-A axis (*r* = 0.31, *p*_spin_ = 0.035; Fig. 8f) and the largest effects were enriched at the S-A axis’s association pole (fifth quintile enrichment analysis: *p*_spin_ = 0.031; Fig. 8g). Together, these findings highlight the generalizability of our developmental and environmental results linking thalamocortical structural connectivity to the S-A axis.

## Discussion

During early stages of cortical neurodevelopment, thalamocortical axons exert powerful control over the arealization, lamination, and neurobiological specification of cortical areas^20–23,25^. In this work, we tested the hypothesis that the thalamus also influences child and adolescent windows of cortical plasticity and provide evidence of synchronized maturation between the cortex and thalamocortical connectivity. To overcome known challenges of thalamic tractography, we first created and anatomically validated a new high-resolution diffusion atlas composed of cortically-annotated thalamocortical structural connections. Applying this atlas to data from three youth cohorts, we reproducibly demonstrated that the development of thalamocortical connectivity is globally heterogeneous, temporally hierarchical, and spatially aligned with variability in cortical developmental profiles. Specifically, we showed that the maturation of thalamocortical structural pathways progresses along the S-A axis and parallels the development of putative non-invasive readouts of cortical developmental plasticity. In this maturational scheme, thalamocortical pathways that mature latest connect to transmodal association regions that are integral to cognitive control and that show signatures indicative of protracted circuit malleability. We furthermore defined relationships between thalamocortical connectivity and neighborhood environment conditions that increase in strength along the S-A axis, thus aligning with the dominant mode of brain-environment associations observed in the cortex during youth^2^. Together, these results uncover that thalamocortical connections develop in a hierarchical and environment-dependent manner across the cortex, consistent with a role for the thalamus in orchestrating the S-A axis of child and adolescent cortical development.

Mechanistic studies in animal models have shown that the thalamus influences the temporal unfolding of developmental processes throughout the span of cortical ontogeny. In early embryonic development, thalamic axons influence the speed of cortical progenitor cell proliferation by releasing a diffusible factor that affects cell cycle kinetics^46^. As development progresses, the rate at which thalamocortical axons grow determines the temporal emergence of regionally-specific cortical maps (e.g., somatotopic maps)^47^. During early postnatal development, experience-dependent transfer of homeoproteins from the thalamus to cortical PV interneurons impacts the timing of sensory cortex critical periods^48^. As maturation decelerates, the stabilization of thalamocortical synapses onto PV interneurons helps to terminate periods of developmental plasticity^8,11^. Animal studies thus point to the thalamus as a timekeeper of cortical neurodevelopment. In the present work, we extend this line of research to humans by linking the timing of thalamocortical connection maturation to the hierarchical progression of maturational processes along the human cortex.

The thalamus’s central position in global modes of brain connectivity and communication^18,49^ makes it well-suited to be a subcortical regulator of experience-dependent cortical development. The thalamus is richly interconnected with a diverse mosaic of cortical regions^16–18^ as well as with sensory systems that process the external world, allowing it to serve as a nexus that can link evolving developmental schedules to physical, cognitive, and social environmental demands. The thalamus has also been identified as a central “gate” that supports indirect cortico-cortical activity propagation, specifically gating information transfer up hierarchically organized processing streams^18,27,29,49^. Though originally identified for cortical communication over short timescales, this thalamic gate may operate developmentally to propagate maturational sequences along the S-A axis. Early in development, exposure to new environmental inputs elicits a marked change in activity in primary cortices that is relayed through thalamic axons and that initiates activity-dependent cortical remodeling. As primary sensory and motor circuits are structurally refined, there is a gradual shift in their functional architecture and the patterning of their intrinsic activity^2,43,44^. Speculatively, this stereotyped refinement of intrinsic activity that occurs during cortical maturation may alter functional signals relayed back to the thalamus via cortical-thalamic projections. This feedback could lead to a consequent shift in thalamic inputs to higher-order cortex that initiates activity-dependent plasticity at the next stage of the processing hierarchy. In this system, cortical activity motifs communicated to the thalamus would provide a biological readout of maturational state, and the thalamus serves as a gate that propagates developmental sequences up the hierarchical S-A axis.

Understanding how the brain regulates hierarchical trajectories of plasticity should facilitate the identification of biological factors that influence windows of environmental sensitivity. We therefore examined associations between thalamocortical connectivity and multiple features of the childhood environment, observing distributed associations between connection properties and neighborhood (but not household) socioeconomic conditions. Notably, associations with neighborhood environment conditions were not homogeneous across the brain. Environment effects were relatively larger for thalamocortical connections that matured for longer, consistent with an accumulation of environmental influences enabled by protracted developmental malleability. As a result, environment effects increased in strength for thalamic connections to regions ranked highest in the S-A axis—mirroring the S-A patterning of associations between neighborhood conditions and regional activity previously described in the cortex^2^. Similarly patterned expression of environmental influences on thalamocortical and cortical properties may indicate that thalamic signals promote environment-dependent sculpting of the cortex in youth. More broadly, these environmental findings add to behavioral observations that neighborhood living conditions can impact long-term outcomes through developmental pathways^50^. These findings furthermore suggest that environmental impacts on the brain continue to accrue throughout development, emphasizing how interventions aimed at mitigating exposures to disadvantaged environments in adolescence could still help support the health of the developing brain.

The present *in vivo* analysis of thalamocortical structural connections in youth has several important limitations. First, white matter pathways identified with diffusion tractography are non-directional, thus the connections studied here inherently contain inputs from thalamus to cortex as well as from cortex to thalamus. Causal investigations in animal models will be needed to study the isolated effects of thalamic projections to cortex on the timing of cortical development. Second, diffusion MRI and FA do not directly measure axonal pathways but rather aggregate directions of local water diffusion. Thus, a percentage of signal ascribed to thalamocortical structural connections may be influenced by diffusion induced by nearby connections in the same voxel. Third, we aimed to study relationships between age-related change in thalamocortical connections and cortical properties by comparing group-level developmental hallmarks. Future work delineating within-individual longitudinal relationships between the development of thalamocortical projections and cortical properties will help to probe these relationships at the individual level. Fourth, we studied associations between thalamocortical connectivity and neighborhood environment factor scores that robustly summarize many interrelated features of the environment. This approach precludes inference regarding which environmental features (e.g., access to material resources, cognitive enrichment, pollution, toxins) or associated psychosocial stressors or protective factors contribute to the associations observed here. Additional studies designed to parse which facets of the environment sculpt brain development and plasticity will be important for informing environmental policies that support youth living across socioeconomic circumstances.

The maturation of the human cerebral cortex follows spatiotemporally precise sequences during its prolonged neurodevelopmental course. The current study bridges animal findings and reproducible developmental neuroimaging to provide evidence linking the thalamus to the dominant sequence of child and adolescent cortical development. It furthermore identifies thalamocortical connectivity as an environmentally responsive biological system that could influence individual differences in the extent and timing of cortical developmental plasticity–and thus windows of developmental vulnerability and opportunity. Further insight into how thalamocortical connectivity affects individualized trajectories of cortical development could ultimately help to inform interventions that reduce the burden of psychopathology in youth by promoting their capacity for adaptive cortical malleability.

## Methods

### Creation of a diffusion atlas of thalamocortical connections

Our analytic approach began with the creation of a population-level tractography atlas comprised of spatially-specific connections between the thalamus and ipsilateral regions of cortex. This population-level atlas was instrumental for the subsequent reliable delineation of regionally-specific thalamocortical pathways in data from individual participants. To create this atlas, we used a publicly available (https://brain.labsolver.org/hcp_template.html) population-average diffusion MRI template that was constructed using data from 1,065 participants included in the HCP Young Adult cohort^51^ (1200-subject release, ages 22-37 years, 575 females). The construction of this diffusion template has been described in detail^52,53^. Briefly, high-resolution, multi-shell diffusion MRI scans were acquired from the 1,065 participants at b-values of 1,000, 2,000, and 3,000 s/mm^2^ (90 directions per shell) and with 1.25 mm isotropic voxels. Preprocessed data were reconstructed using q-space diffeomorphic reconstruction (QSDR)^54^, which performs generalized q-sampling imaging (GQI) in MNI ICBM152 2009a space. GQI is a model-free diffusion MRI reconstruction approach that estimates diffusion orientation distribution functions (ODFs) directly from the diffusion signal (the distribution of diffusion spins) to resolve complex fiber orientations^55^. GQI-based ODFs can be estimated in single-shell and multi-shell diffusion sampling schemes. QSDR outputs were aggregated across the 1,065 participants to build a population-averaged template of voxelwise diffusion distributions. We used this QSDR diffusion template in DSI Studio for construction of the thalamocortical connectivity atlas.

We first used DSI Studio to perform deterministic fiber tracking on the QSDR diffusion template to identify 2 million streamlines terminating in the left thalamus and 2 million streamlines terminating in the right thalamus. Contralateral white matter masks were used as regions of avoidance during hemisphere-specific thalamic tracking to only extract streamlines between the thalamus and ipsilateral brain regions. Deterministic tractography in DSI Studio uses voxel ODFs and quantitative anisotropy measures to resolve crossing fibers, reduce partial volume effects, filter noisy fibers, and define better tracking termination locations^56^. In the context of the present study, deterministic tracking offers advantages for identifying cortically-constrained thalamocortical connections with high validity and high termination specificity (as compared to probabilistic tracking approaches which tend to achieve broader coverage of connection profiles with a trade-off of more false positives and lower endpoint specificity^57–59^). Reflecting this advantage, in an international tractography challenge, the deterministic tractography approach implemented here reconstructed 92% validly connecting streamlines (compared to an average of 54% valid streamlines across all submissions) and additionally identified the lowest number of invalid white matter bundles^60^.

The following parameters were used for thalamic tractography, which were chosen following parameter testing: threshold index = qa, otsu threshold = 0.45, minimum streamline length = 10 mm, maximum streamline length = 300 mm. In addition to these stable parameters, random parameter saturation was used to select the anisotropy threshold, turning angle, step size, and smoothing level for each generated streamline. Random parameter saturation entails using a random combination of the aforementioned tracking parameters (each within a pre-defined, set range of appropriate values) to generate a broad array of viable streamlines. By sampling across the entire appropriate parameter space rather than arbitrarily selecting a single value in the space, this approach can resolve streamlines with varied properties and enhances both reconstruction accuracy and computational reproducibility^61^. Following identification of 2 million streamlines with endpoints in the left and right thalamus, we extracted ipsilateral connections between the thalamus and individual cortical regions defined by the HCP-MMP atlas^62^, which was included with DSI Studio. These connections served as the basis of the thalamocortical tractography atlas.

All regionally-specific thalamocortical connections extracted by the above procedure underwent a process of visual quality assurance and manual curation prior to their inclusion in the final atlas. The macroscale anatomy of extracted connections was compared to prior thalamocortical tractography results and tract tracing accounts, when available, to confirm anatomical accuracy. All connections were additionally subject to potential manual editing to delete false positive streamlines with biologically impractical architectures as well as superfluous streamlines that diverged from the core connection. The manual checking process was also used to entirely remove n = 16 thalamocortical connections from the final atlas that were deemed overly sparse based on the template tracking or that could not be reconstructed in participant-level data in both PNC and HCPD datasets. After curating all extracted thalamocortical connections, skeletonized versions of the final set of connections were generated by deleting “repeat” streamlines with redundant trajectories, operationalized here as streamlines within a distance of < 3 voxels. Removal of redundant streamlines enhances the computational efficiency of the subsequent automated tracking procedure, without compromising anatomical representation or coverage. Skeletonized thalamocortical connections were combined to create a new atlas of human thalamocortical connections (Fig. 1). This atlas was used as a custom atlas with DSI Studio’s automated tractography (replacing the built-in ICBM152_adult atlas) to study thalamocortical connection maturation in developmental datasets.

### Developmental datasets

Developmental analyses were conducted independently in three large, cross-sectional youth datasets: PNC, HCPD, and HBN. PNC and HCPD were used as the primary discovery and replication datasets for all study analyses. HBN, a sample of youth presenting with psychiatric concerns, was additionally included to assess whether key developmental and environmental findings replicated in a clinical sample. All subsequent methods concern these three datasets. In all three studies (PNC, HCPD, HBN), all participants over the age of 18 years gave informed consent prior to participating in the study. Participants under the age 18 gave informed assent and were enrolled with written consent from their legal guardians. Participants in all three studies received monetary compensation for participation; HBN participants additionally received diagnostic evaluations and referral information.

PNC study procedures were approved by the Institutional Review Boards of both the University of Pennsylvania and the Children’s Hospital of Philadelphia. HCPD study procedures were approved by a central Institutional Review Board at Washington University in St. Louis. HBN study procedures were approved by the Chesapeake Institutional Reviewer Board.

#### PNC

PNC^63^ is a community sample of children and adolescents from a broad range of socioeconomic circumstances that were residing in the greater Philadelphia area (Pennsylvania, USA). Initial exclusion criteria for the PNC were minimal and included inability to engage in psychiatric and cognitive phenotyping, impaired vision or hearing, and medical problems that could impact brain function (described in detail in Satterthwaite et al., 2016). Data from 1,145 PNC participants were included in the current study. Participants ranged in age from 8 to 23 years (mean age of 15.3 ± 3.5 years) and had a sex distribution of 608 females and 537 males (sex was self-reported; intersex was not assessed). Participants self-identified their race and ethnicity: 0.3% of participants identified as American Indian or Alaskan Native, 0.9% as Asian, 42.4% as Black or African American, 10.5% as multiracial, and 45.9% as White.

#### HCPD

HCPD^64^ is a sample of children and adolescents that were recruited at four academic sites including the University of Minnesota (Minnesota, USA), Harvard University (Massachusetts, USA), Washington University in St. Louis (Missouri, USA) and University of California-Los Angeles (California, USA). HCPD was designed to be a US population-representative study of typical brain development that included participants from varied geographical, ethnic, and socioeconomic backgrounds. Initial exclusion criteria for this study are detailed in a prior report^64^; notable exclusions included premature birth, serious neurological and endocrine conditions, requiring special services at school, treatment for a psychiatric illness for > 12 months, and hospitalization for a psychiatric condition for > 2 days. Data from 572 HCPD participants were included in the current study (lifespan 2.0 release). Participants ranged in age from 8 to 22 years (mean age of 14.8 ± 3.9 years) and had a sex distribution of 306 females and 266 males (sex was self-reported; intersex was not assessed). Participants self-identified their race and ethnicity: 7.7% of participants identified as Asian, 10.3% as Black or African American, 14.0% as multiracial, and 65.2% as White (data were missing for 2.8% of participants).

#### HBN

HBN^65^ is a sample of children and adolescents from the New York City area (New York, USA) that were referred to the study due to concerns about psychiatric symptoms. The HBN initiative was created by the Child Mind Institute to sample a broad range of commonly encountered forms of clinically-significant youth psychopathology; as a result, study exclusions were generally minimal^5^. Extensive information about the HBN sample is provided in Alexander et al., 2017^5^. Approximately 85% of participants in this sample meet criteria for a clinical disorder, including but not limited to anxiety, depressive, attention deficit and hyperactivity, conduct, impulse control, autism spectrum, learning, communication, and stress disorders. Data from 959 HBN participants were included in the current study. Participants ranged in age from 8 to 22 years (mean age of 12.2 ± 3.1 years) and had a sex distribution of 329 females and 585 males (sex was self-reported; intersex was not assessed; 45 participants were missing a binary sex indication and were assigned a sex of other). Participants self-identified their race and ethnicity: 2.5% of participants identified as Asian, 13.5% as Black or African American, 46.5% as White, and 18.2% as not belonging to these three race categories (data were missing for 19.3% of participants).

### MRI Acquisition

This study uses T1-weighted (T1w) structural images, diffusion-weighted images, and field maps collected from the three developmental datasets.

#### PNC

MRI data collected from all PNC participants were acquired on the same 3T Siemens TIM Trio Scanner at the University of Pennsylvania. The T1w images were acquired with a magnetization-prepared rapid acquisition gradient-echo (MPRAGE) sequence with the following parameters: repetition time of 1,810 ms, echo time of 3.51 ms, inversion time of 1,100 ms, flip angle of 9 degrees, 160 slices, and a voxel resolution of 0.94 × 0.94 × 1 mm. Single shell diffusion scans were acquired with a b-value = 1,000 s/mm^2^ in 64 directions with 7 interspersed scans with b = 0 s/mm^2;^ all volumes were acquired in the anterior-posterior direction. The collection of these 71 volumes was divided between two runs. The following parameters were used for the diffusion acquisition: repetition time of 8,100 ms, echo time of 82 ms, and a voxel resolution of 1.875 x 1.875 x 2 mm. In addition to the structural and diffusion acquisitions, a phase-difference based field map was acquired to facilitate susceptibility distortion correction of the diffusion data. Field maps were acquired with a double-echo, gradient-recalled echo (GRE) sequence with a repetition time of 1,000 ms, echo times of 2.69 ms and 5.27 ms; a flip angle of 60 degrees, 44 slices, and a voxel resolution of 3.75 x 3.75 x 4 mm.

#### HCPD

HCPD MRI scans were acquired at 4 sites on 3T Siemens Prisma scanners (MR derivatives were harmonized across sites as described below). T1w images were acquired with a 3D multi-echo MPRAGE sequence with an in-plane acceleration factor of 2 and the following additional parameters: repetition time of 2,500 ms, echo times of 1.8, 3.6, 5.4, and 7.2 ms, inversion time of 1,000 ms, flip angle of 8 degrees, 208 slices, and a voxel resolution of 0.8 mm isotropic. Diffusion scans were acquired in four consecutive runs with two shells of b = 1,500 and 3,000 s/mm^2^ with a multiband factor of four. 92-93 directions were acquired per shell (370 directions total) along with 28 total b = 0 s/mm^2^ volumes for a total of 398 volumes. Of the 370 diffusion-weighted volumes, 185 distinct directions were each acquired twice with opposite phase encoding directions (anterior-posterior and posterior-anterior). The following parameters were used for the diffusion acquisition: repetition time of 3,230 ms, echo time of 89 ms, and a voxel resolution of 1.5 mm isotropic.

#### HBN

This study analyzes 3T MRI data from the HBN, which were acquired at 3 sites (MR derivatives were harmonized across sites as described below). Data were collected at the Rutgers University Brain Imaging Center on a 3T Siemens Tim Trio scanner as well as at the CitiGroup Cornell Brain Imaging Center and the CUNY Advanced Science Research Center on 3T Siemens Prisma Scanners. T1w images were acquired with a MPRAGE sequence with the following parameters: repetition time of 2,500 ms, echo time of 3.15 ms, inversion time of 1,060 ms, flip angle of 8 degrees, 224 slices, and a voxel resolution of 0.8 mm isotropic. Diffusion scans were acquired with a multiband factor of three in two shells of b = 1,000 and 2,000 s/mm^2^ in the anterior-posterior phase encoding direction. 64 directions were acquired per shell (128 directions total) along with 1 b = 0 volume. The following parameters were used for the diffusion acquisition: repetition time of 3,320 ms, echo time of 100.2 ms, and a voxel resolution of 1.8 mm isotropic. A reverse phase encoding b = 0 was additionally acquired for use as an EPI-based field map in susceptibility distortion correction.

### Diffusion MRI preprocessing

Diffusion scans (and corresponding T1w images and fieldmaps) from PNC, HCPD, and HBN were preprocessed using QSIPrep^66^, which is based on Nipype and uses Nilearn, Dipy, ANTs, FSL, and software tools described below. QSIPrep versions 0.14.2 and 0.16.1 were used in PNC/HBN and HCPD, respectively. The same sequence of preprocessing steps were performed in both datasets. However, given that slightly different QSIPrep versions were applied to PNC/HBN and HCPD, the software versions used by its internal operations differed. The following internal software versions were used in PNC/HBN and HCPD, respectively: Nipype 1.6.1 and 1.8.5, Nilearn 0.8.0 and 0.9.2, ANTs 2.3.1 and 2.4.0, FSL 6.0.3 and 6.0.5.

Anatomical processing was also performed within QSIPrep. T1w images were corrected for intensity non-uniformity using N4BiasFieldCorrection^67^ (ANTs), skull-stripped using antsBrainExtraction (ANTs) with OASIS as a target template, and spatially normalized to the nonlinear ICBM152 2009c template using nonlinear registration with antsRegistration. For diffusion data processing, a series of preprocessing steps were applied separately to data from each diffusion run (2 runs in PNC; 4 runs in HCPD; 1 run in HBN) and then runs were concatenated. Any images with a b-values < 100 s/mm^2^ were considered b = 0 volumes. First, MP-PCA denoising, as implemented in dwidenoise (MRtrix3^68^), was applied with a 5-voxel window. After MP-PCA, Gibbs unringing (MRtrix3) was performed with mrdegibbs. Following unringing, B1 field inhomogeneity was corrected using dwibiascorrect (MRtrix3), which implements the N4 algorithm^67^. After B1 bias correction, the mean intensity of the diffusion-weighted series was adjusted so the mean intensity of all b = 0 images matched across separate runs. QSIPrep was additionally used to correct for head motion, eddy currents, and susceptibility distortions. FSL’s eddy was used for head motion and eddy current correction^69^. In all datasets, eddy was configured with a q-space smoothing factor of 10, a total of 5 iterations, and 1000 voxels used to estimate hyperparameters. Eddy’s outlier replacement was run^70^.

Given that PNC, HCPD, and HBN acquired different versions of fieldmaps (GRE and EPI), different approaches were taken to correct for susceptibility artifacts. In the PNC, fieldmap-based susceptibility distortion correction was performed by creating a B0 map using a phase-difference image and a magnitude image from the GRE fieldmap acquisition. In HCPD and HBN, reverse phase-encoding EPI-based fieldmaps were collected, resulting in pairs of images with distortions going in opposite directions. For EPI-based susceptibility distortion correction, b = 0 references images with reversed phase encoding directions were used along with an equal number of b = 0 images extracted from diffusion scans. From these pairs the susceptibility-induced off-resonance field was estimated. Fieldmaps were ultimately incorporated into the eddy current and head motion correction interpolation. Final interpolation was performed using the Jacobian modulation (jac) method. After preprocessing, the concatenated diffusion data were resampled to AC-PC to be in alignment with the T1w data. The output resolution of the fully preprocessed diffusion data was 1.8 mm isotropic in PNC and HBN and 1.5 mm isotropic in HCPD.

### Sample construction

PNC, HCPD, and HBN participants with T1w images, field maps, and identical-parameter^71^ diffusion MRI scans (i.e., non-variant acquisitions) available were considered for inclusion in the present work. Exclusion criteria were applied to available scans in each dataset to ensure that only high quality neuroimaging data from youth without serious medical conditions were analyzed. In all three datasets, participants were excluded from the initial neuroimaging sample for medical problems that could impact brain function or incidentally encountered abnormalities in their neuroimaging data (medical exclusion). Participants were additionally excluded if they had low quality T1w images with motion, artifacts, or poor surface reconstructions (T1w quality exclusion), if their raw diffusion scans were missing gradient directions (diffusion directions exclusion), if in-scan head motion during the diffusion scan exceeded a mean framewise displacement of 1 mm (diffusion motion exclusion), or if their preprocessed diffusion scans had a low neighborhood correlation (diffusion quality exclusion). The neighborhood correlation metric quantifies the average pairwise spatial correlation between pairs of diffusion volumes that sample similar points in q-space, with lower correlations reflecting reduced data quality. The range of neighborhood correlation values obtained will vary by the diffusion scan acquisition parameters and noise level, thus dataset-specific exclusion thresholds were used (< 0.9 in PNC, < 0.6 in HCPD, < 0.7 in HBN).

PNC was used as the discovery dataset in this study. In PNC, 1,358 individuals had all required neuroimaging data available. From this original neuroimaging sample, 213 total participants (15.7%) were ultimately excluded for the above criteria (applied successively) including: n = 118 for medical exclusions, n = 25 for the T1w quality exclusion, n = 57 for the diffusion quality exclusion, and a remaining n = 13 for the diffusion motion exclusion. HCPD was used as the main replication dataset. In HCPD, 640 individuals had neuroimaging data available. From this originally eligible sample, 54 total participants (10.6%) were excluded for the above criterion including: n = 7 for medical exclusions, n = 8 for the diffusion directions exclusion, n = 15 for the diffusion quality exclusion, and then n = 24 for the diffusion motion exclusion. In addition, in order to directly compare developmental results from HCPD to PNC, children less than 8 years old were excluded from HCPD to match the PNC age range (n = 14; only 2.4% of the remaining eligible sample). This young age exclusion was additionally important in HCPD as there is likely insufficient data available from individuals < 8 years old for accurate developmental modeling in this dataset (the same age range exclusion has been applied in the past by independent groups for this reason^6^). HBN was used as a secondary replication dataset to assess if findings could be extended to a predominantly psychiatric sample. As with HCPD, we only considered data from participants that overlapped with the PNC age range. 1,530 participants from data releases 1-9 in HBN had non-variant (collected with the same acquisition) T1w and diffusion scans available. In this initial sample, 1,149 were between the ages of 8 and 22 years. From the 1,149 individuals, 190 total participants (16.5%) were ultimately excluded for the above criterion including: n = 53 for the T1w quality exclusion, n = 10 for the diffusion quality exclusion, and then n = 127 for the diffusion motion exclusion.

### Delineation of individual-specific thalamocortical pathways

In order to consistently identify the same thalamocortical connections across PNC, HCPD, and HBN participants, we used our population-level thalamocortical tractography atlas as an anatomical prior with DSI Studio’s trajectory-based automated tract recognition (auto-track) approach^53,61^. To conduct auto-track, we first reconstructed participants’ preprocessed diffusion scans with GQI. GQI was executed using QSIPrep’s GQI reconstruction workflow (version 0.16.0RC3 in all datasets) with a ratio of mean diffusion distance of 1.25. Auto-track uses a distance-based pathway recognition approach to best match streamlines generated in individual participant’s data to connections included in the thalamocortical connectivity atlas. Specifically, the auto-track procedure involves non-linearly warping thalamocortical atlas connections to participant diffusion data, seeding deterministic tractography in voxels that correspond to each connection in the atlas, comparing generated streamlines to atlas connections using the Hausdorff distance, and discarding streamlines with a Hausdorff distance above a selected threshold (i.e., those with trajectories that diverge from the atlas connection). We used a relatively strict Hausdorff distance threshold of 10 to ensure identification of connections with similar shapes and anatomical endpoints across individuals while still allowing for subtle person-specific differences in anatomy. The following specific parameters were used for deterministic tracking: threshold index = qa, otsu threshold = 0.50, a track:voxel ratio of 4, a yield rate of .0000001, and no topology-informed pruning. As with the tractography atlas, random parameter saturation was used to select the anisotropy threshold, turning angle, step size, and smoothing level for each generated streamline.

Nearly all thalamocortical structural connections represented in the population-level atlas could be clearly identified in diffusion data from all youth participants. Few connections, however, were either not identified or only sparsely reconstructed in participant data. These unsuccessful reconstructions at the participant-level provide important evidence that the thalamocortical tractography pipeline employed here was not over-fitting participants’ data to identify connections that were not faithfully represented in the diffusion signal. To ensure that only well-represented thalamocortical connections were analyzed in the present work, we applied exclusion criterion to connections at individual and dataset levels. In particular, for each participant, we excluded connections that had < 5 streamlines from consideration. After applying this participant-level exclusion, we then only analyzed connections for which at least 90% of participants had a valid (>= 5 streamlines) connection. As a result, 15 connections (6.3%) present in the original thalamocortical connectivity atlas were not studied in PNC, 8 connections (3.4%) in HCPD, and 13 connections (5.5%) in HBN.

### Derivation of thalamocortical connectivity properties

For all reconstructed thalamocortical connections, we computed metrics that provide information about connection microstructure and about where the connection’s streamlines terminated within the thalamus. Specifically, we calculated two metrics, FA and thalamic C-M_t_ gradient position, in all analyzed connections from all youth datasets. FA is a diffusion tensor-derived measure sensitive to tissue microstructure that has been studied in the developmental literature, has good test-retest reliability, and can be appropriately calculated from both single and multi-shell acquisitions. We used DSI Studio to compute the average FA along each thalamocortical connection. FA was calculated based on the single-shell of b = 1,000 s/mm^2^ in PNC and shells < 1,750 s/mm^2^ in HCPD and HBN. High b-value shells (> 1,750) were not used for the calculation of FA as high b-values sensitive to non-gaussian diffusion do not meet the assumptions of the tensor fit, and can introduce noise or inaccuracies into DTI values. Note, however, that all shells were used to enhance tractography performance. Given that HCPD and HBN diffusion data were collected across different scanners, we used ComBat to harmonize thalamocortical FA measures within each dataset to mitigate scanner-specific effects. ComBat has previously been shown to remove unwanted scanner effects from FA measures when age is the biological phenotype of interest, and to do so more effectively than alternate harmonization approaches^72^. ComBat was implemented via the neuroCombat package in R (https://github.com/Jfortin1/neuroCombat_Rpackage). Age was protected as a biological variable of interest in ComBat along with developmental model covariates of sex and in-scan head motion.

We additionally derived a C-M_t_ value for every delineated thalamocortical connection, which quantifies where the thalamic endpoints of the connection fall in a within-thalamus core-matrix gradient. To accomplish this, we used the core-matrix thalamic gradient^30^ from Müller et al., 2020 (obtained from https://github.com/macshine/corematrix in MNI space). This thalamic gradient was produced based on the relative mRNA expression of parvalbumin (PVALB) and calbindin (CALB1) genes, which are markers for core and matrix thalamic cell types, respectively. We calculated the average C-M_t_ value of all voxels that corresponded to a given structural connections’ thalamic termination area by transforming a mask of each thalamocortical connection to MNI space using warps computed during preprocessing in QSIPrep. The mean C-M_t_ value of all thalamic voxels within the connection’s mask was then computed.

### General approach to hypothesis testing

Throughout this work, we evaluate the statistical significance of GAM-derived smooth terms and linear terms as well as the significance of spatial correlations between brain maps. Here we note general approaches used in significance testing, which will be described in further detail below. When assessing significance with GAMs, which were applied individually to each thalamocortical connection, the false discovery rate correction was used to correct p-values (*p*_FDR_) for multiple comparisons across all connections within a dataset and statistical significance was set at *p*_FDR_ < 0.05. When correlating brain maps, we used non-parametric Spearman’s rank-based correlations throughout to assess monotonic relationships between variables, except when comparing the same metric derived independently in two datasets (e.g., age effects in PNC and HPCD) as linear relationships were expected. To compute the significance of each correlation comparing two brain maps, we employed spin-based, spatial rotation tests or “spin tests”^73^. Spin tests compute a p-value (*p*_spin_) by comparing the empirically observed correlation between two brain maps to a null distribution of correlations obtained by spatially rotating (spinning) spherical projections of the brain maps that maintain their native spatial covariance structure. The *p*_spin_ value is computed as the number of times the rotation-based correlation value is greater than (for positive empirical correlations) or less than (for negative empirical correlations) the empirical correlation value, divided by the total number of spatial rotations (spins). We used a null of 10,000 spatial rotations throughout. To carry out spin tests with thalamocortical connectivity data, we projected connection-derived metrics to the cortical surface for spinning by assigning each HCP-MMP cortical region the value of its corresponding thalamic connection. Region-based spin tests were implemented using the rotate_parcellation algorithm in R (https://github.com/frantisekvasa/rotate_parcellation). Spin tests were additionally used to create a null distribution of values for anatomical and environment enrichment tests (described further below). All statistics were performed in R version 4.2.3. All statistical tests were two-sided. Statistical analyses were conducted separately in each of the three developmental datasets.

### Maps of cortical organization, function, and development

We integrated brain maps of cortical feature variability, functional diversity, and developmental heterochronicity into the present study to provide insights into thalamocortical connection anatomy and development. To facilitate this, the HCP-MMP atlas was used to parcellate data from the brain maps described below.

#### The sensorimotor-association axis of cortical organization

Every HCP-MMP region was assigned a specific rank in the S-A axis based on its position in this dominant motif of cortical feature organization. S-A axis ranks were previously computed in Sydnor et al., 2021^1^ by averaging rank orderings of ten cortical feature maps that exhibit systematic and concerted spatial variation between primary sensory and motor cortices and transmodal association cortices. These maps include the principal gradient of functional connectivity^74^, the cortical T1/T2 ratio^75^, macroscale MRI measurements of cortical thickness, allometric scaling calculated as local area scaling with changes in total brain size^76^, metabolic demand indexed by aerobic glycolysis^77^, cerebral perfusion estimated by arterial spin labeling^78^, functional diversity determined by spatial variation in Neurosynth meta-analytic decodings, cytoarchitectural similarity measured using the BigBrain atlas^79^, a dominant mode of gene expression proxied by the first principal component of brain-expressed genes^75^, and macaque-to-human evolutionary cortical expansion^80^. Accordingly, the S-A axis represents a large-scale, prominent axis of cortical organization that captures the stereotyped patterning of numerous macrostructural, microstructural, functional, metabolic, transcriptomic, and evolutionary features across the human cerebral cortex. The rank ordering of all HCP-MMP regions along the S-A axis was obtained from https://github.com/PennLINC/S-A_ArchetypalAxis. Low ranking regions at the axis’s sensorimotor pole are nearly exclusively primary and early unimodal cortices. Mid-ranking regions span from multimodal cortices involved in multisensory integration and language to those involved in attention and working memory. Higher ranking regions are involved in decision making capacities. Regions at the axis’s association pole are strongly involved in cognitive control, social cognition, self-referential thought, and emotion regulation.

#### Neurosynth meta-analytic association maps

Neurosynth^81^ version 0.7 was used to create HCP-MMP meta-analytic maps for a diversity of psychological terms capturing both lower-order and higher-order psychological processes. In accordance with prior work^82^ we obtained meta-analytic activation maps for 123 cognitive terms that were present in both the Neurosynth database and the Cognitive Atlas^83^; maps were obtained using the Neuroimaging Meta-Analysis Research Environment (NiMARE)^84^. Term-specific maps were computed in volumetric space using multilevel kernel density Chi-square analysis, mapped to the fslr surface, and parcellated with the HCP-MMP parcellation. Values in term-specific maps are association test *z*-scores that quantify the extent to which activation in a cortical region occurred more consistently in prior functional MRI studies that mentioned a given term as compared to studies that did not.

#### Brain charts of developmental heterochronicity

We used previously published maps characterizing across-cortex variability in the development of the E/I ratio, myelin, and intrinsic activity amplitude as indexed by non-invasive MRI measures sensitive to these cortical properties. The cortical E/I ratio was non-invasively estimated by Zhang, Larsen et al. 2023^5^ by fitting a biophysically plausible circuit model to resting state functional MRI data from youth in the PNC (*N* = 855, ages 8-23 years). The magnitude of maturational decline in the E/I ratio was quantified across the cortex through a linear regression between model-estimated E/I ratio and age, producing an age slope map that was parcellated with the HCP-MMP atlas. Age-related change in cortical myelin content was studied by Baum et al., 2022^6^ by calculating the T1/T2 ratio in data from HCPD participants (*N* = 628, ages 8-22 years). These authors determined the annualized rate of cortical myelin growth in each HCP-MMP region by fitting GAMs with a smooth function for age and quantifying the difference between the model-estimated T1/T2 ratio at ages 21 and 8, divided by the number of years in this range (a point estimate for the rate was estimated by posterior sampling). Intrinsic activity fluctuation amplitude was studied in Sydnor et al., 2023^2^ using resting-state functional MRI data from the PNC sample (*N* = 1,033, ages 8-23 years). Fluctuation amplitude was quantified as the average power of low frequency functional MRI fluctuations, given that increases in power in the frequency domain are mathematically proportional to increases in signal amplitude in the time domain. The age at which fluctuation amplitude began to decrease was evaluated in every cortical region by fitting GAMs with thin plate regression splines to model developmental trajectories and identifying the youngest age at which the first derivative of the developmental spline was significantly negative.

### Analysis of thalamocortical connectivity atlas characteristics

After creating a diffusion atlas of human thalamocortical connections, we assessed whether cortical regions without connections included in the atlas (“absent”) had smaller surface areas and greater sulcal depth than regions with thalamic connections represented in the atlas (“present”). To statistically test this, we computed the average surface area of cortical regions with absent thalamic connections, and compared this empirical value to a null distribution of surface areas obtained by spatially rotating the cortical surface area map. A spin testing-based *p*_spin_ value was computed by calculating the number of times the empirical surface area value was less than the null value. The same spin testing procedure was implemented to determine whether regions with absent connections had significantly greater sulcal depth than those with present connections.

### Analysis of thalamocortical connection anatomical properties

To study whether thalamocortical connections exhibited different patterns of connectivity with sensory versus association cortices, we assessed whether thalamocortical connection FA and C-M_t_ values exhibited ordered variation along the S-A axis. As described above, FA and C-M_t_ values were calculated for every thalamocortical connection reconstructed in every individual. To study alignment of these connectivity measures with the S-A axis, the average (across-participant) value was computed for each connection. Spearman’s correlations between connection FA or C-M_t_ values and S-A axis ranks were then calculated and the significance of these correlations were determined with spin tests.

### Analysis of thalamocortical connection development

#### Developmental modeling

To quantitatively characterize diversity in the development of structural connections between the thalamus and individual cortical regions, we fit connection-specific GAMs with FA as the dependent variable, age as a smooth term, and sex and diffusion scan head motion (mean framewise displacement) as linear covariates. Models were fit separately for each individual thalamocortical connection using thin plate regression splines as the smooth term basis set and the restricted maximal likelihood approach for selecting smoothing parameters. All models were fit using the mgcv package^85^ in R. We used GAMs for developmental modeling given that they are capable of capturing a broad array of both linear and non-linear age-dependent relationships and can furthermore be harnessed to characterize age windows of significant developmental change and change offset. Each GAM produces a smooth function for age that is generated from a linear combination of weighted basis functions (splines) and that represents the connection’s developmental trajectory. As in our prior work^2^, to prevent overfitting of the flexible smooth function we set the maximum basis complexity (k) to 3. A value of k = 3 was selected given that model fits were relatively non-complex (requiring a smaller number of knots) and the k-index indicated this basis complexity was sufficient. Higher values of k were moreover visually deemed to result in overfitting in multiple connections.

After fitting connection-specific GAMs, we interrogated GAM-derived properties including the significance of the age effect, the magnitude of the age effect, and the age at which the connection stopped exhibiting significant developmental change. The significance of the age effect, which denotes the significance of the smooth function quantifying the relationship between FA and age, was determined by an ANOVA F-test that compared the full GAM model to a nested, reduced model with no age term (in the form of a generalized likelihood ratio test). A significant F-test (FDR-corrected; *p*_FDR_) indicates that including the smooth term for age in the model significantly improved the model fit as compared to a model with only sex and diffusion scan head motion as predictors. The magnitude of the age effect was calculated as the partial *R*^2^ between the full GAM model and the reduced model without age. To resolve the age at which each connection stopped exhibiting significant developmental change during childhood and adolescence—which can be interpreted as the connection’s age of maturation—we first calculated the first derivative of the age smooth function using finite differences, which provides an age-specific rate of developmental change (Δ FA / Δ age). We then identified the youngest age at which the rate of developmental change was not significantly different from 0 by obtaining a simultaneous 95% confidence interval for these first derivatives and pinpointing when the confidence interval first included 0. Derivative analyses utilized the gratia package^86^ in R. After fitting main developmental GAMs, we additionally fit models with a sex by age (factor-smooth) interaction to assess whether developmental trajectories of FA significantly differed by sex. The significance of the interaction term was extracted and corrected for multiple comparisons (*p*FDR).

#### Developmental characterization

A primary goal of this research was to empirically evaluate the hypothesis that the development of thalamocortical structural connections is coupled to the heterochronous patterning of developmental change across the cortex’s S-A axis. We assessed this hypothesis by characterizing how differences in ages of thalamocortical connection maturation were related to (1) cortical function, (2) cortical and thalamic axes, and (3) cortical developmental heterogeneity. To facilitate these three analyses, we computed the age of maturation for each connection and projected these ages onto HCP-MMP regions.

First, to functionally decode cortical regions with earlier and later maturing thalamic connections, we computed Spearman’s correlations between the cortex-projected thalamocortical age of FA maturation map and each of the 123 psychological term meta-analytic maps made using Neurosynth. In this analysis, psychological terms with the most negative correlation values exhibit high fMRI activation-based z-scores in cortical regions with the earliest maturing thalamic connections. Terms with the most positive correlations display highest z-scores in cortical regions with the latest maturing thalamic connections. To provide insight into the types of functions ascribed to regions with early versus late maturing thalamocortical connections, we therefore identified the 10 most negatively correlated and 10 most positively correlated cognitive terms that emerged in a given dataset (thus extracting 20 developmentally-pertinent terms total). We also constructed a term-overlap null distribution to statistically assess correspondence of the 20 terms across independent developmental datasets. For this term-overlap null, we computed the number of overlapping terms (n/20) obtained between two datasets as well as the probability of obtaining >= n overlapping terms when randomly drawing 20 terms per dataset from the full 123 term list (without replacement; 10,000 iterations). This null distribution of term overlaps was used to calculate a permutation-based *p*_PERM_ value.

Second, to further characterize how differences in thalamocortical connection development were instantiated across the cortex and thalamus, we correlated thalamocortical connection maturational ages with the S-A axis, with the cortex’s three primary anatomical axes (A-P, D-V, and M-L axes), and with the core-matrix thalamic gradient using Spearman’s correlations. We ascertained the significance (FDR-corrected) of these individual correlations with spin tests. We additionally assessed whether the correlation with the S-A axis was significantly greater in magnitude than correlations with A-P, D-V, and M-L cortical axes and the core-matrix gradient by implementing statistical tests (FDR-corrected) for comparing two dependent, overlapping correlations. These tests used a back-transformed average Fisher’s *Z* procedure to compare correlations and were executed using the cocor package^87^ in R with the hittner2003 test.

Third, we aimed to study correspondence between the spatiotemporal patterning of maturational variability across cortical regions and across thalamocortical structural connections. To test for correspondence between cortical and thalamocortical development, we used the three previously described cortical charts of developmental heterochronicity. Specifically, we used Spearman’s correlations and spin tests to relate the thalamocortical age of maturation map to developmental maps of biophysical model-estimated E/I ratio, T1/T2 ratio-indexed cortical myelin, and functional MRI-based fluctuation amplitude.

### Analysis of environmental effects

#### Measures of youths’ socioeconomic environments

We examined relationships between thalamocortical connection FA and socioeconomic features of youths’ household environments (education, income) and neighborhood environments (geocoding-derived factor scores). Not all datasets collected the same environmental information, thus the most relevant variables available were studied here. Household socioeconomic position was proxied by the average years of education obtained by both (when data was available) or one caregiver in both PNC and HCPD. Caregiver years of education were directly reported in PNC and were missing for 8 participants who were excluded from this analysis. In HCPD, caregiver education information was reported as the highest educational level achieved and was therefore recoded to a numerical value to allow for continuous analyses. Grade levels were directly recoded (e.g., “8^th^ grade” to 8 years, “10^th^ grade” to 10 years) and degrees were recoded based on average years to degree completion in the United States (e.g., “Bachelor’s degree” to 16 years, “Professional degree” to 19 years, “Doctoral degree” to 22 years). Household socioeconomic position was additionally estimated by the income-to-needs ratio in HCPD (income was not provided in PNC). This ratio was calculated by dividing total annual family income by the federal poverty line based on the participant’s total family size. A city cost of living-adjusted ratio was then computed to account for geographic area (St. Louis = 1, Minneapolis = 1.175, Los Angeles = 1.558, Boston = 1.624). To mitigate the potential impact of extreme income outliers and account for the right-skewed distribution of these data, the city-adjusted income-to-needs ratio was winsorized (lowest and highest 1% of the data) and the natural logarithm was taken, in accordance with prior studies^37,88^.

Information about the neighborhood, rather than household, environment was obtained using geocoding and was not available in HCPD given that participant addresses were not released. In PNC, data about each individual’s neighborhood (block-level) environment were extracted using home addresses and the census-based American Community Survey. Census-based variables were factor analyzed in Moore et al., 2016^89^ to derive a factor score for each participant that captured multivariate features of the neighborhood environment a child lived in. The first factor from an exploratory two-factor analysis performed in Moore et al., 2016 was used here, which has the following factor loadings: the percentage of residents who were married (loading = 0.85), median family income (0.82), the percentage of residents with a high school education (0.74), the percentage of residents who were employed (0.68), median age (0.61), the percentage of residents who were female (−0.26), the percentage of houses that were vacant (−0.60), population density (−0.71), and the percentage of residents in poverty (−0.86).

Although neighborhood environment information could not be obtained in HCPD, participant addresses were collected in HBN, allowing us to assess whether neighborhood environment results replicated in a second sample. Mirroring the approach taken in the PNC, a neighborhood environment factor score was calculated for HBN participants by factor analyzing geocoded variables at the level of census block-groups. Specifically, variables from the American Community Survey and Environmental Protections Agency were factor analyzed using exploratory structural equation modeling, and five factors were extracted. The neighborhood socioeconomic factor included here had the following loadings: median family income (loading = 0.89), median home value (0.80), median owner upkeep cost (0.80), median rent (0.78), the percent of residents with a high school education (0.57), median rooms per dwelling (0.52), the percentage of residents who were married (0.50), the percent of residents without health insurance (-0.49), and the percent of residents in poverty (-0.61). Address information and associated neighborhood environment scores were missing for 13 individuals in HBN who were excluded from this analysis.

#### Statistical testing of environmental associations

To model relationships between thalamocortical connection FA and environmental features, we re-fit developmental GAMs with environment variables included as linear continuous covariates. Model terms included a smooth term for age and linear terms for sex, diffusion scan head motion, and the environment variable under study (caregiver education, income, or neighborhood factor score). This model formulation allowed us to identify age-independent main effects of the environment across developmental stages. Significance of the linear environment term was FDR corrected (*p*_FDR_) across all connection-specific GAMs in an analysis. The direction and magnitude of the relationship between connection FA and the environment variable was determined by the *t* value of the linear environment term, which we refer to as the statistical environment effect. In addition to the above GAMs used to quantify main effects of the environment, we furthermore fit GAMs with an age by environment varying coefficient interaction to model how associations between the environment variable and connection FA vary over the smooth function of age. These varying coefficient interaction GAMs were used to predict connection-specific FA developmental trajectories for varying levels of the environment variable.

#### Environmental characterization

In a final series of analyses, we explored how significant environment effects varied across the S-A axis during youth. These analyses were only conducted for neighborhood environment effects, given that caregiver education and income effects were ultimately not significant. We first conducted an effect enrichment analysis wherein we tested whether the across-connection distribution of environment effects (*t* values) was homogeneous, or whether effects were significantly smaller or larger in magnitude for connections to certain portions of the S-A axis. To execute this enrichment analysis, we first computed the average environment *t* value in five quintiles of the S-A axis. We then compared this empirical *t* to a null distribution of quintile-specific *t* values obtained by applying 10,000 spherical spatial rotations to the cortex-projected environment effect map, producing a spin-based enrichment *p*_spin_. To complement the quintile enrichment analysis, we furthermore correlated significant environment effects with the S-A axis and with A-P, D-V, and M-L cortical axes and the core-matrix thalamic gradient using Spearman’s correlations (with FDR-corrected spin tests). We compared the magnitude of correlations between the S-A axis and each of these four axes using the previously described cocor test (FDR-corrected) for comparing two dependent, overlapping correlations.

## Data availability

The present work utilized existing developmental data from the PNC (https://www.ncbi.nlm.nih.gov/projects/gap/cgi-bin/study.cgi?study_id=phs000607.v3.p2), the Lifespan 2.0 HCPD release (https://nda.nih.gov/ccf), and the Healthy Brain Network (https://fcon_1000.projects.nitrc.org/indi/cmi_healthy_brain_network/). These data are publicly available for download from the provided links. This study furthermore incorporated a diffusion MRI QSDR template (1.25 mm population-averaged FIB file) constructed from HCP Young Adult data that was made available (by F-C Yeh) at https://brain.labsolver.org/hcp_template.html. Analyses presented here used multiple cortical and thalamic annotations including the thalamic core-matrix gradient (distributed in MNI space at https://github.com/macshine/corematrix), Neurosynth meta-analytic maps (made available through NiMARE at https://nimare.readthedocs.io/en/stable/), and the archetypal S-A axis (accessible via https://pennlinc.github.io/S-A_ArchetypalAxis/). The maps of T1/T2 ratio development (https://balsa.wustl.edu/study/P2DmK) and fluctuation amplitude development (https://github.com/PennLINC/spatiotemp_dev_plasticity/blob/main/cortical_maps/AgeofDeclineOnset _FirstNegDeriv.pscalar.nii) are also publicly accessible via the accompanying links. The thalamocortical connectivity atlas created here from diffusion MRI is available for download along with instructions for employing it with DSI Studio at https://github.com/PennLINC/thalamocortical_development/tree/main/results/thalamocortical_autotrac k_template.

## Code availability

Neuroimaging data were processed with containerized software packages. Diffusion MRI data were preprocessed and reconstructed with qsiprep (https://hub.docker.com/r/pennbbl/qsiprep/tags). Thalamocortical tractography was produced for the population-level atlas and for individual participants using a containerized version of DSI studio https://hub.docker.com/r/dsistudio/dsistudio (version chen-2023-02-17). All additional study analyses and statistics were conducted in bash, python, and R using original analysis code. All study code is provided at https://github.com/PennLINC/thalamocortical_development/tree/main. A detailed guide to code implementation and study replication describing all analytic steps can be accessed at https://pennlinc.github.io/thalamocortical_development/.

## Acknowledgements

This research was supported by the National Institutes of Health, including by grants T32MH016804 to V.J.S.; R00MH127293 to B.L.; R01MH132934 and R01MH133843 to A.F.A-B.; R01MH124045, P50MH109429, R01MH133334, R01MH130578 and R01MH131638 to A.R.F and M.P.M.; R01MH119219 to R.E.G and R.C.G; T32DC000038 and F31HD111139 to S.L.M; R01MH120174 and R01MH119185 to D.R.R.; R01MH123550, R01MD123563, and R01NS112274 to R.T.S; R01MH113550 to T.D.S and D.S.B; R01MH120482 to T.D.S and M.P.M; R01MH112847 to T.D.S and R.T.S.; R37MH125829 to T.D.S.; and R01EB022572 to T.D.S Research was also supported by a National Science Foundation Graduate Research Fellowship (DGE-1845298 to V.J.S.) and by the CHOP Research Institute, the Penn-CHOP Lifespan Brain Institute, the Penn AI2D center, and the AE Foundation.

## Author Contributions

V.J.S. conceived of and designed the study. B.L., M.C., and T.D.S. provided scientific and technical mentorship throughout the study lifecycle. D.M.B., A.R.F., R.E.G., R.C.G., M.P.M., and T.D.S. provided resources and supervised collection of the developmental neuroimaging data. V.J.S., S.C., K.M., S.L.M., D.R.R., T.S., and M.C. curated and processed the developmental neuroimaging data using software developed by S.C., K.M., T.S., F-C.Y., and M.C. T.M.M and G.S. generated the neighborhood environment factor scores. J.S. provided data used to derive the S-A axis. E.J.M. and J.M.S. derived the core-matrix thalamic gradient. V.J.S. created and curated the thalamocortical tractography atlas with technical support from P.A.C., F-C.Y. and M.C. V.J.S. implemented all statistical analyses with guidance from B.L., D.S.B., A.F.A-B., R.T.S., and T.D.S. M.J.A. provided expert input on thalamocortical connectivity anatomy. D.M.B and A.P.M provided expert input on environmental analyses. V.J.S. wrote all study-specific code and created all figures. J.B. and M.C. conducted an internal code review and technical replication of all study findings. V.J.S wrote the original manuscript. All authors reviewed and revised the manuscript.

## Competing Interests

The authors declare no competing interests.

## Materials and Correspondence

Please address correspondence and material requests to authors Valerie J. Sydnor and Theodore D. Satterthwaite.

## Notes

### Competing Interest Statement

The authors have declared no competing interest.

## References

1. Sydnor, V. J. et al. Neurodevelopment of the association cortices: Patterns, mechanisms, and implications for psychopathology. Neuron 109, 2820–2846 (2021).

2. Sydnor, V. J. et al. Intrinsic activity development unfolds along a sensorimotor–association cortical axis in youth. Nat Neurosci 26, 638–649 (2023).

3. Larsen, B. & Luna, B. Adolescence as a neurobiological critical period for the development of higher-order cognition. Neurosci Biobehav Rev 94, 179–195 (2018).

4. Larsen, B. et al. A developmental reduction of the excitation:inhibition ratio in association cortex during adolescence. Sci Adv 8, eabj8750 (2022).

5. Zhang, S. et al. In vivo whole-cortex marker of excitation-inhibition ratio indexes cortical maturation and cognitive ability in youth. Proceedings of the National Academy of Sciences 121, e2318641121 (2024).

6. Baum, G. L. et al. Graded Variation in T1w/T2w Ratio during Adolescence: Measurement, Caveats, and Implications for Development of Cortical Myelin. J. Neurosci. 42, 5681–5694 (2022).

7. Benoit, L. J. et al. Adolescent thalamic inhibition leads to long-lasting impairments in prefrontal cortex function. Nat Neurosci 25, 714–725 (2022).

8. Ribic, A. & Biederer, T. Emerging Roles of Synapse Organizers in the Regulation of Critical Periods. Neural Plast 2019, 1538137 (2019).

9. Coleman, J. E. et al. Rapid Structural Remodeling of Thalamocortical Synapses Parallels Experience-Dependent Functional Plasticity in Mouse Primary Visual Cortex. J Neurosci 30, 9670–9682 (2010).

10. Faini, G. et al. Perineuronal nets control visual input via thalamic recruitment of cortical PV interneurons. eLife 7, e41520 (2018).

11. Ribic, A., Crair, M. C. & Biederer, T. Synapse-Selective Control of Cortical Maturation and Plasticity by Parvalbumin-Autonomous Action of SynCAM 1. Cell Reports 26, 381–393.e6 (2019).

12. Quast, K. B. et al. Rapid synaptic and gamma rhythm signature of mouse critical period plasticity. Proceedings of the National Academy of Sciences 120, e2123182120 (2023).

13. Larsen, B., Sydnor, V. J., Keller, A. S., Yeo, B. T. T. & Satterthwaite, T. D. A critical period plasticity framework for the sensorimotor–association axis of cortical neurodevelopment. Trends in Neurosciences 46, 847–862 (2023).

14. Takesian, A. E. & Hensch, T. K. Chapter 1 - Balancing Plasticity/Stability Across Brain Development. in Progress in Brain Research (eds. Merzenich, M. M., Nahum, M. & Van Vleet, T. M.) vol. 207 3–34 (Elsevier, 2013).

15. Reh, R. K. et al. Critical period regulation across multiple timescales. Proc Natl Acad Sci U S A 117, 23242–23251 (2020).

16. Howell, A. M. et al. The spatial extent of anatomical connections within the thalamus varies across the cortical hierarchy in humans and macaques. eLife 13, (2024).

17. Oldham, S. & Ball, G. A phylogenetically-conserved axis of thalamocortical connectivity in the human brain. Nat Commun 14, 6032 (2023).

18. Sherman, S. M. & Guillery, R. W. Distinct functions for direct and transthalamic corticocortical connections. Journal of Neurophysiology 106, 1068–1077 (2011).

19. Alzu’bi, A., Homman-Ludiye, J., Bourne, J. A. & Clowry, G. J. Thalamocortical Afferents Innervate the Cortical Subplate much Earlier in Development in Primate than in Rodent. Cerebral Cortex 29, 1706–1718 (2019).

20. Antón-Bolaños, N., Espinosa, A. & López-Bendito, G. Developmental interactions between thalamus and cortex: a true love reciprocal story. Curr Opin Neurobiol 52, 33–41 (2018).

21. O’Leary, D. D. M., Chou, S.-J. & Sahara, S. Area Patterning of the Mammalian Cortex. Neuron 56, 252–269 (2007).

22. Vue, T. Y. et al. Thalamic Control of Neocortical Area Formation in Mice. J. Neurosci. 33, 8442– 8453 (2013).

23. Li, H. et al. Laminar and columnar development of barrel cortex relies on thalamocortical neurotransmission. Neuron 79, 970–986 (2013).

24. Murakami, T., Matsui, T., Uemura, M. & Ohki, K. Modular strategy for development of the hierarchical visual network in mice. Nature 608, 578–585 (2022).

25. Shibata, M. et al. Regulation of prefrontal patterning and connectivity by retinoic acid. Nature 598, 483–488 (2021).

26. Park, S. et al. A shifting role of thalamocortical connectivity in the emergence of cortical functional organization. Nat Neurosci 1–11 (2024) doi:10.1038/s41593-024-01679-3.

27. Theyel, B. B., Llano, D. A. & Sherman, S. M. The corticothalamocortical circuit drives higher-order cortex in the mouse. Nat Neurosci 13, 84–88 (2010).

28. Jaramillo, J., Mejias, J. F. & Wang, X.-J. Engagement of Pulvino-cortical Feedforward and Feedback Pathways in Cognitive Computations. Neuron 101, 321–336.e9 (2019).

29. Sherman, S. M. Thalamus plays a central role in ongoing cortical functioning. Nat Neurosci 19, 533–541 (2016).

30. Müller, E. J. et al. Core and matrix thalamic sub-populations relate to spatio-temporal cortical connectivity gradients. Neuroimage 222, 117224 (2020).

31. Erisir, A. & Dreusicke, M. Quantitative morphology and postsynaptic targets of thalamocortical axons in critical period and adult ferret visual cortex. Journal of Comparative Neurology 485, 11– 31 (2005).

32. Cruikshank, S. J., Lewis, T. J. & Connors, B. W. Synaptic basis for intense thalamocortical activation of feedforward inhibitory cells in neocortex. Nat Neurosci 10, 462–468 (2007).

33. Fagiolini, M. & Hensch, T. K. Inhibitory threshold for critical-period activation in primary visual cortex. Nature 404, 183–186 (2000).

34. Fair, D. A. et al. Maturing Thalamocortical Functional Connectivity Across Development. Front Syst Neurosci 4, 10 (2010).

35. Avery, S. N. et al. Development of Thalamocortical Structural Connectivity in Typically Developing and Psychosis Spectrum Youths. Biological Psychiatry: Cognitive Neuroscience and Neuroimaging 7, 782–792 (2022).

36. Tooley, U. A. et al. Prenatal environment is associated with the pace of cortical network development over the first three years of life. 2023.08.18.552639 Preprint at 10.1101/2023.08.18.552639 (2023).

37. Noble, K. G. et al. Family income, parental education and brain structure in children and adolescents. Nature Neuroscience 18, 773–778 (2015).

38. Sun, C. et al. Human Thalamic-Prefrontal Peduncle Connectivity Revealed by Diffusion Spectrum Imaging Fiber Tracking. Frontiers in Neuroanatomy 12, (2018).

39. Liu, M., Lerma-Usabiaga, G., Clascá, F. & Paz-Alonso, P. M. Reproducible protocol to obtain and measure first-order relay human thalamic white-matter tracts. NeuroImage 262, 119558 (2022).

40. Wiesendanger, R. & Wiesendanger, M. The thalamic connections with medial area 6 (supplementary motor cortex) in the monkey (macaca fascicularis). Exp Brain Res 59, 91–104 (1985).

41. Jones, E. G. The thalamic matrix and thalamocortical synchrony. Trends Neurosci 24, 595–601 (2001).

42. McGee, A. W., Yang, Y., Fischer, Q. S., Daw, N. W. & Strittmatter, S. M. Experience-Driven Plasticity of Visual Cortex Limited by Myelin and Nogo Receptor. Science 309, 2222–2226 (2005).

43. Golshani, P. et al. Internally mediated developmental desynchronization of neocortical network activity. J Neurosci 29, 10890–10899 (2009).

44. Wu, M. W., Kourdougli, N. & Portera-Cailliau, C. Network state transitions during cortical development. Nat. Rev. Neurosci. 1–18 (2024) doi:10.1038/s41583-024-00824-y.

45. Michael, C. et al. Socioeconomic resources in youth are linked to divergent patterns of network integration and segregation across the brain’s transmodal axis. bioRxiv 2023.11.08.565517 (2023) doi:10.1101/2023.11.08.565517.

46. Dehay, C., Savatier, P., Cortay, V. & Kennedy, H. Cell-Cycle Kinetics of Neocortical Precursors Are Influenced by Embryonic Thalamic Axons. J Neurosci 21, 201–214 (2001).

47. Fetter-Pruneda, I. et al. Shifts in Developmental Timing, and Not Increased Levels of Experience-Dependent Neuronal Activity, Promote Barrel Expansion in the Primary Somatosensory Cortex of Rats Enucleated at Birth. PLOS ONE 8, e54940 (2013).

48. Sugiyama, S. et al. Experience-dependent transfer of Otx2 homeoprotein into the visual cortex activates postnatal plasticity. Cell 134, 508–520 (2008).

49. Shine, J. M., Lewis, L. D., Garrett, D. D. & Hwang, K. The impact of the human thalamus on brain-wide information processing. Nat Rev Neurosci 24, 416–430 (2023).

50. Chetty, R., Hendren, N. & Katz, L. F. The Effects of Exposure to Better Neighborhoods on Children: New Evidence from the Moving to Opportunity Experiment. American Economic Review 106, 855–902 (2016).

51. Glasser, M. F. et al. The Human Connectome Project’s neuroimaging approach. Nat Neurosci 19, 1175–1187 (2016).

52. Yeh, F.-C. et al. Population-Averaged Atlas of the Macroscale Human Structural Connectome and Its Network Topology. Neuroimage 178, 57–68 (2018).

53. Yeh, F.-C. Population-based tract-to-region connectome of the human brain and its hierarchical topology. Nat Commun 13, 4933 (2022).

54. Yeh, F.-C. & Tseng, W.-Y. I. NTU-90: a high angular resolution brain atlas constructed by q-space diffeomorphic reconstruction. Neuroimage 58, 91–99 (2011).

55. Yeh, F.-C., Wedeen, V. J. & Tseng, W.-Y. I. Generalized q-Sampling Imaging. IEEE Transactions on Medical Imaging 29, 1626–1635 (2010).

56. Yeh, F.-C., Verstynen, T. D., Wang, Y., Fernández-Miranda, J. C. & Tseng, W.-Y. I. Deterministic Diffusion Fiber Tracking Improved by Quantitative Anisotropy. PLOS ONE 8, e80713 (2013).

57. Schilling, K. G. et al. Tractography dissection variability: What happens when 42 groups dissect 14 white matter bundles on the same dataset? NeuroImage 243, 118502 (2021).

58. Sarwar, T., Ramamohanarao, K. & Zalesky, A. Mapping connectomes with diffusion MRI: deterministic or probabilistic tractography? Magnetic Resonance in Medicine 81, 1368–1384 (2019).

59. Côté, M.-A. et al. Tractometer: Towards validation of tractography pipelines. Medical Image Analysis 17, 844–857 (2013).

60. Maier-Hein, K. H. et al. The challenge of mapping the human connectome based on diffusion tractography. Nature Communications 8, 1349 (2017).

61. Yeh, F.-C. Shape analysis of the human association pathways. Neuroimage 223, 117329 (2020).

62. Glasser, M. F. et al. A multi-modal parcellation of human cerebral cortex. Nature 536, 171–178 (2016).

63. Satterthwaite, T. D. et al. The Philadelphia Neurodevelopmental Cohort: A publicly available resource for the study of normal and abnormal brain development in youth. Neuroimage 124, 1115–1119 (2016).

64. Somerville, L. H. et al. The Lifespan Human Connectome Project in Development: A large-scale study of brain connectivity development in 5–21 year olds. NeuroImage 183, 456–468 (2018).

65. Alexander, L. M. et al. An open resource for transdiagnostic research in pediatric mental health and learning disorders. Scientific Data 4, 1–26 (2017).

66. Cieslak, M. et al. QSIPrep: an integrative platform for preprocessing and reconstructing diffusion MRI data. Nat Methods 18, 775–778 (2021).

67. Tustison, N. J. et al. N4ITK: Improved N3 Bias Correction. IEEE Trans Med Imaging 29, 1310– 1320 (2010).

68. Tournier, J.-D. et al. MRtrix3: A fast, flexible and open software framework for medical image processing and visualisation. NeuroImage 202, 116137 (2019).

69. Andersson, J. L. R. & Sotiropoulos, S. N. An integrated approach to correction for off-resonance effects and subject movement in diffusion MR imaging. NeuroImage 125, 1063–1078 (2016).

70. Andersson, J. L. R., Graham, M. S., Zsoldos, E. & Sotiropoulos, S. N. Incorporating outlier detection and replacement into a non-parametric framework for movement and distortion correction of diffusion MR images. Neuroimage 141, 556–572 (2016).

71. Covitz, S. et al. Curation of BIDS (CuBIDS): A workflow and software package for streamlining reproducible curation of large BIDS datasets. Neuroimage 263, 119609 (2022).

72. Fortin, J.-P. et al. Harmonization of multi-site diffusion tensor imaging data. Neuroimage 161, 149–170 (2017).

73. Alexander-Bloch, A. F. et al. On testing for spatial correspondence between maps of human brain structure and function. Neuroimage 178, 540–551 (2018).

74. Margulies, D. S. et al. Situating the default-mode network along a principal gradient of macroscale cortical organization. Proc Natl Acad Sci U S A 113, 12574–12579 (2016).

75. Burt, J. B. et al. Hierarchy of transcriptomic specialization across human cortex captured by structural neuroimaging topography. Nature Neuroscience 21, 1251 (2018).

76. Reardon, P. K. et al. Normative brain size variation and brain shape diversity in humans. Science 360, 1222–1227 (2018).

77. Vaishnavi, S. N. et al. Regional aerobic glycolysis in the human brain. Proc Natl Acad Sci U S A 107, 17757–17762 (2010).

78. Satterthwaite, T. D. et al. Impact of puberty on the evolution of cerebral perfusion during adolescence. Proc Natl Acad Sci U S A 111, 8643–8648 (2014).

79. Paquola, C. et al. The BigBrainWarp toolbox for integration of BigBrain 3D histology with multimodal neuroimaging. eLife 10, e70119 (2021).

80. Hill, J. et al. Similar patterns of cortical expansion during human development and evolution. Proc Natl Acad Sci U S A 107, 13135–13140 (2010).

81. Yarkoni, T., Poldrack, R. A., Nichols, T. E., Van Essen, D. C. & Wager, T. D. Large-scale automated synthesis of human functional neuroimaging data. Nat Methods 8, 665–670 (2011).

82. Shafiei, G. et al. Topographic gradients of intrinsic dynamics across neocortex. eLife 9, e62116 (2020).

83. Poldrack, R. et al. The Cognitive Atlas: Toward a Knowledge Foundation for Cognitive Neuroscience. Frontiers in Neuroinformatics 5, (2011).

84. Salo, T. et al. NiMARE: Neuroimaging Meta-Analysis Research Environment. Aperture Neuro 3, 1–32 (2023).

85. Wood, S. N. *Generalized Additive Models: An Introduction with R*. (Chapman and Hall/CRC, New York, 2017). doi:10.1201/9781315370279.

86. Gavin L. Simpson. gratia: Graceful ggplot-Based Graphics and Other Functions for GAMs Fitted using mgcv. https://gavinsimpson.github.io/gratia/ (2022).

87. Diedenhofen, B. & Musch, J. cocor: A Comprehensive Solution for the Statistical Comparison of Correlations. PLoS One 10, e0121945 (2015).

88. Weissman, D. G. et al. Family income is not significantly associated with T1w/T2w ratio in the Human Connectome Project in Development. Imaging Neuroscience 1, 1–10 (2023).

89. Moore, T. M. et al. Characterizing social environment’s association with neurocognition using census and crime data linked to the Philadelphia Neurodevelopmental Cohort. Psychological Medicine 46, 599–610 (2016).

